# Cyclin D3 drives inertial cell cycling in dark zone germinal center B cells

**DOI:** 10.1101/2020.11.17.385716

**Authors:** Juhee Pae, Jonatan Ersching, Tiago B. R. Castro, Marta Schips, Luka Mesin, Samuel J. Allon, Jose Ordovas-Montanes, Coraline Mlynarczyk, Ari Melnick, Alejo Efeyan, Alex K. Shalek, Michael Meyer-Hermann, Gabriel D. Victora

## Abstract

During affinity maturation, germinal center (GC) B cells alternate between proliferation and somatic hypermutation in the dark zone (DZ) and affinity-dependent selection in the light zone (LZ). This anatomical segregation imposes that the vigorous proliferation that allows clonal expansion of positively-selected GC B cells takes place ostensibly in the absence of the signals that triggered selection in the LZ, as if by “inertia.” We find that such inertial cycles specifically require the cell cycle regulator cyclin D3. Cyclin D3 dose-dependently controls the extent to which B cells proliferate in the DZ and is essential for effective clonal expansion of GC B cells in response to strong T follicular helper (Tfh) cell help. Introduction into the *Ccnd3* gene of a Burkitt lymphoma-associated gain-of-function mutation (T283A) leads to larger GCs with increased DZ proliferation and, in older mice, to clonal B cell lymphoproliferation, suggesting that the DZ inertial cell cycle program can be coopted by B cells undergoing malignant transformation.

## Introduction

Germinal centers (GCs) are the sites of affinity maturation, the process by which antibodies improve their affinity for antigen over time (Cyster and Allen, 2019; De Silva and Klein, 2015; Eisen, 2014; Mesin et al., 2016; Rajewsky, 1996; Shlomchik et al., 2019; Victora and Nussenzweig, 2012). For efficient affinity maturation, GC B cells must cycle between two major transcriptional states, associated with localization of B cells to each of the two microanatomical “zones” of the GC. When in the dark zone (DZ),B cells proliferate vigorously while they mutate their immunoglobulin genes by somatic hypermutation (SHM). After transition to the light zone (LZ),B cells bearing advantageous mutations are selectively driven to clonally expand, based at least in part on their ability to bind and present antigen to GC-resident T follicular helper (Tfh) cells (Victora et al., 2010). Successive cycles of somatic hypermutation and affinity-based selection ultimately enrich for higher-affinity cells in among the GC B cell population, in a process known as cyclic re-entry (MacLennan, 1994; Victora and Nussenzweig, 2012).

A unique consequence of the anatomical compartmentalization of the GC is that mitogenic signals are segregated from the proliferation they induce. Upon positive selection, B cells typically transition from G1 to S phase of the cell cycle in the LZ, migrate from LZ to DZ while in S phase, and undergo G2 and M phases in the DZ (Gitlin et al., 2014; Victora et al., 2010). After this first division, cell cycling continues in the DZ, with most B cells undergoing on average two additional cell cycles before returning to the LZ for further selection (Gitlin et al., 2014). GC B cells that receive stronger signals from Tfh cells in the LZ, however, can undergo a much greater number of proliferative cycles in the DZ, resulting in exponential clonal expansion (Gitlin et al., 2014; Meyer-Hermann et al., 2012; Victora et al., 2010). At its extreme, this regulated expansion can lead to “clonal bursts,” where a single B cell can take over a 2,000-cell GC in the course of a few days (Tas et al., 2016). These bursts are associated with massive diversification of generally higher-affinity SHM variants, and as such are likely to play an important role in the generation of high-affinity B cell clones (Amitai et al., 2017; Bannard and Cyster, 2017; Mesin et al., 2016).

Despite the importance of DZ proliferation for GC B cell selection and affinity maturation, our understanding of GC B cell selection has historically focused on events that take place in the LZ (Mesin et al., 2016; Shlomchik et al., 2019). Consequently, the precise mechanisms that allow DZ proliferation to take place in the apparent absence of direct mitogenic signals are still incompletely understood. For instance, upon interaction with Tfh cells, positively-selected LZ B cells express the transcription factor c-Myc (Calado et al., 2012; Dominguez-Sola et al., 2012; Finkin et al., 2019), as well as the mammalian target of rapamycin (mTOR)-mediated anabolic program (Ersching et al., 2017). While both c-Myc and mTOR are required for LZ B cells to migrate to the DZ and enter the proliferative phase, the induction and effect of these pathways appear to be mostly restricted to the LZ (Dominguez-Sola et al., 2012; Ersching et al., 2017). This suggests that DZ B cells retain a memory of the intensity of the c-Myc and mTORC1-dependent “charge” they received previously in the LZ, and that they subsequently translate this memory into the number of cell cycles they will undergo in the DZ using a cell-intrinsic “timer” or “counter” (Bannard et al., 2013; Gitlin et al., 2014). Whereas the transcriptional effects c-Myc can be partly extended to the DZ through engagement of the AP-4 transcription factor (Chou et al., 2016), the molecular pathways that directly control the number of cycles a GC B cell undergoes in the DZ remain uncharacterized.

Here, we used a combination of unbiased single-cell RNA-sequencing (scRNA-seq) analysis and targeted genetic and pharmacological manipulation of cell cycle regulators to investigate the molecular nature of DZ cell cycles. We find that DZ B cells adopt a distinct E2Fhigh/c-Myclow mode of cell cycling that allows rapid and continuous proliferation in the absence of external mitogenic signals. We show that the cell cycle regulator cyclin D3, previously shown to be required for GC formation and maintenance (Cato et al., 2011; Peled et al., 2010), is a specific, dose-dependent controller of this phenotype and of DZ cell cycling, loss of which cannot be overcome by LZ cycling induced by strongly increased Tfh cell help. Introduction into mice of a gain-of-function mutation in cyclin D3 derived from human Burkitt lymphoma (Schmitz et al., 2014; Schmitz et al., 2012) leads to exacerbated B cell proliferation specifically in the GC DZ and to development of clonal post-GC B cell expansions, linking the inertial proliferative program to malignant transformation.

## Results

### B cell proliferation in the DZ is Tfh-independent

To better understand GC B cell cycling in the DZ, we first sought to formally determine whether S phase entry by DZ B cells requires acute signals from Tfh cells in addition to those delivered in the LZ. To this end, we used a synchronized selection model (Ersching et al., 2017; Victora et al., 2010) to allow kinetically precise blocking of Tfh cell help to GC B cells at set time points after induced positive selection (Fig. 1A). In this model, we use adoptive transfer of 4-hydroxy-3-nitro-phenylacetyl (NP)-specific B1-8^hi^ B cells followed by immunization with NP conjugated to chicken ovalbumin (NP-OVA) to create GCs in which the majority of B cells lack the surface receptor DEC-205 (encoded by the gene *Ly75*). Treatment of mice with ongoing GCs with an antibody to DEC-205 fused to OVA (DEC-OVA) leads to presentation of OVA peptides by *Ly75*^+/+^ GC B cells only. This induces *Ly75*^+/+^ cells to preferentially interact with Tfh cells, triggering their positive selection (Pasqual et al., 2015; Victora et al., 2010). In this process, GC B cells also become synchronized: 12 h after DEC-OVA, most *Ly75*^+/+^ cells are located in the LZ, where they interact extensively with Tfh cells and induce cell cycle progression (Ersching et al., 2017; Shulman et al., 2014). By 36 h after DEC-OVA, most *Ly75*^+/+^ cells have transitioned to the DZ, where they begin their proliferative burst (Victora et al., 2010). Figures 1A-C and S1 provide an overview of key time points in these kinetics. Entry into S-phase of the cell cycle, revealed by double-pulsing mice with EdU and BrdU nucleotides (Gitlin et al., 2014), was greatly increased in DZ B cells at 36 h post-DEC-OVA, indicating that GC B cells progress through the G1-S checkpoint while in the DZ compartment (Figs. 1B-C and S1E,F). DZ S-phase entry persisted above the steady state level until at least 60 h post-DEC-OVA treatment, when *Ly75*^+/+^ cells remained predominantly in the DZ and continued to expand (Figs. 1B,C and S1B-D).

**Figure 1.**
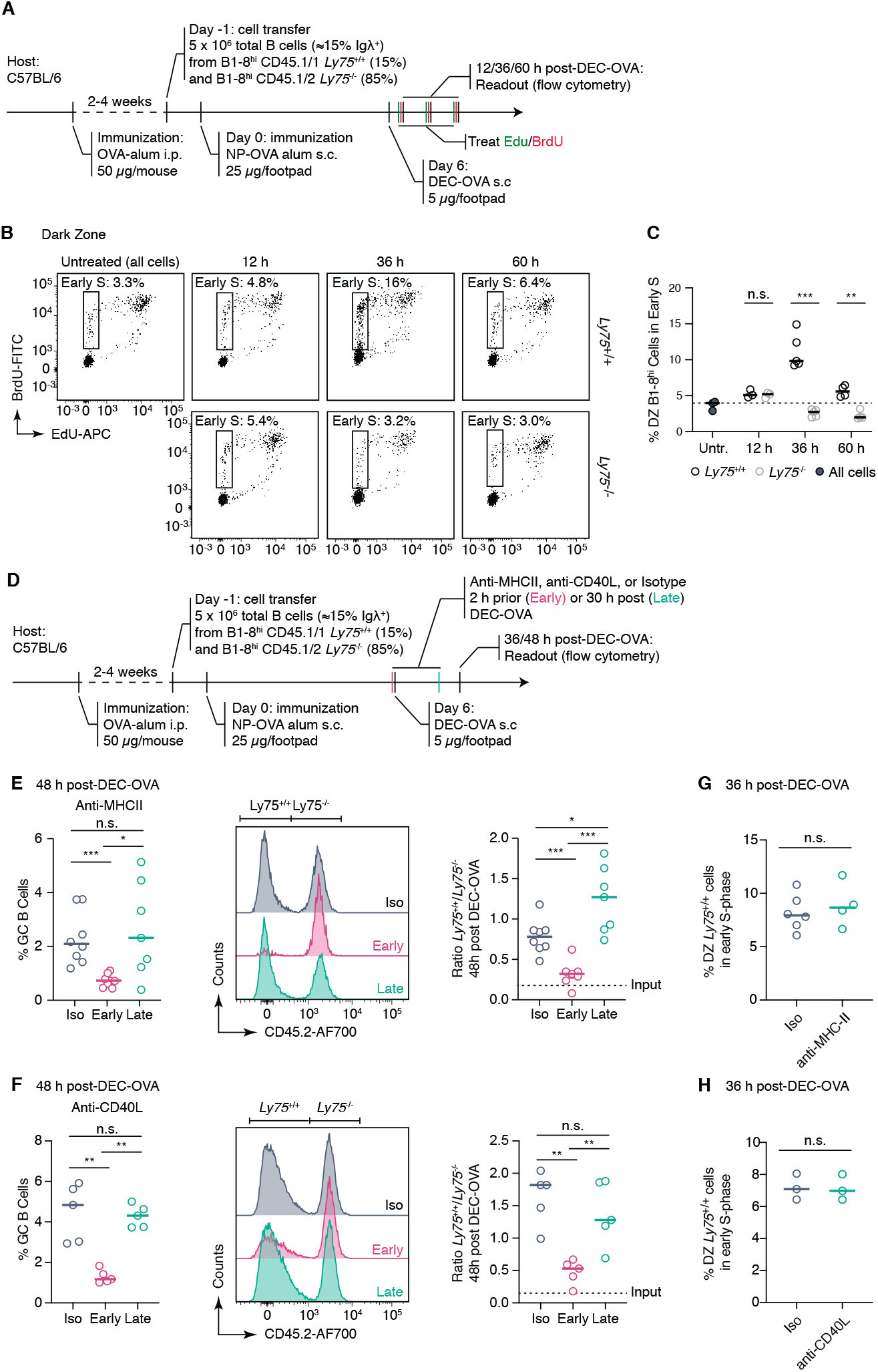
Cell cycle entry by DZ B cells does not require Tfh cell help. **(A)** Experimental setup for DEC-OVA-induced *Ly75*^+/+^ GC B cells followed by double labeling with EdU/BrdU to assay for S phase entry. (B) S phase entry of *Ly75*^+/+^ and *Ly75*^−/−^ B1-8^hi^ cells in the DZ, quantified in (C). **p < 0.01, ***p < 0.001, paired t test comparison between *Ly75*^+/+^ and *Ly75*^−/−^ cells in the same animal. Bars indicate median. Data pooled from two independent experiments. (D) Experimental setup for DEC-OVA-induced positive selection of *Ly75*^+/+^ GC B cells followed by inhibition of T-B interaction using anti-MHCII or anti-CD40L before (early) or after (late) DZ re-entry. (E) Effect on GC size and (F) expansion of *Ly75*^+/+^ cells over *Ly75*^−/−^ upon anti-MHCII treatment. (G) Effect on GC size and (H) expansion of *Ly75*^+/+^ cells over Ly75^−/−^ upon anti-CD40L treatment. *p < 0.05, **p < 0.01, ***p < 0.001, ****p < 0.0001, n.s. non-significant, nonparametric Mann-Whitney test, compared to the group treated with an isotype control. Bars indicate median. Data pooled from three (anti-MHCII) or two (anti-CD40L) independent experiments. (I-J) S phase entry of *Ly75*^+/+^ B1-8^hi^ cells in the DZ upon late treatment with anti-MHCII (I) or anti-CD40L (J). n.s. non-significant, nonparametric Mann-Whitney test, compared to the group treated with an isotype control. Data pooled from two (anti-MHCII) or one (anti-CD40L) independent experiments.

To determine whether T cell help is continually required for DZ proliferation, we acutely blocked Tfh-B cell interaction either at the time of the initial Tfh signal delivery in the LZ (6 h post-DEC-OVA) or following the transition of positively selected B cells to the DZ but prior to the proliferative burst (30 h post-DEC-OVA; Fig. 1D). As expected from the role of Tfh-mediated signaling in this model (Victora et al., 2010), early blocking of T-B interactions in the LZ using antibodies to MHC-II or CD40 ligand (CD40L) effectively prevented clonal expansion of *Ly75*^+/+^ cells by 48 h post-DEC-OVA (Fig. 1E,F). By contrast, blocking either pathway after B cells transitioned to the DZ had little, if any, effect on the expansion of *Ly75*^+/+^ cells over *Ly75*^−/−^ cells (Fig. 1E,F), or on the ability of *Ly75*^+/+^ DZ B cells to enter S-phase, as evidenced by EdU/BrdU incorporation (Fig. 1G,H).We conclude that sustained proliferation of B cells in the GC DZ upon positive selection does not require continuous help from Tfh cells. We therefore refer to DZ cell cycles as “inertial,” since they proceed in a cell-intrinsic fashion according to the strength of the initial “push” from Tfh cells, in contrast to the “reactive” cell cycles that LZ B cells undergo immediately downstream of Tfh-mediated signaling and selection.

### Inertial cycling is sustained by prolonged E2F activation in the absence of c-Myc activity

To identify the transcriptional programs associated with inertial cycling, we performed whole transcriptome single-cell mRNA sequencing (scRNAseq) on GC B cells at different timepoints after forcing positive selection using DEC-OVA. We index-sorted single *Ly75*^+/+^ B cells using antibodies to LZ/DZ markers, and sorted cells were sequenced using the Smart-Seq2 protocol (Trombetta et al., 2014). We assayed GC B cells first at 12 h after DEC-OVA, when *Ly75*^+/+^ B cells are enriched in the LZ in the process of receiving cognate help from Tfh cells, and then at 30, 46, and 60 h after DEC-OVA, as DZ B cells transition from signal-dependent proliferation to predominantly inertial modes of cycling prior to returning to the LZ between 72 and 96 h post-DEC-OVA (Figs. 1A-D and S1) (Victora et al., 2010). For comparison, we also included a sample of *Ly75*^−/−^ counterselected LZ cells from the 12 h timepoint (Supplemental Table 1).

The 1,220 cells that passed our quality control thresholds fell into 7 major clusters (Fig. 2A). These were determined in large part by cell cycle phase as inferred from their transcriptional profile (Tirosh et al., 2016), with a lesser contribution of their LZ/DZ phenotype as defined by surface staining (Fig. 2B). Clusters 0 and 4 contained primarily G1 cells; clusters 2, 3, and 6 were enriched in S-phase cells; and clusters 1 and 5 were enriched in cells in the G2 and M phases (Fig. 2C). To follow the evolution of positively-selected B cells across these clusters as they transitioned from reactive to inertial cell cycles and then to quiescence, we compared *Ly75*^+/+^ and *Ly75*^−/−^ cells in the LZ at 12 h-post DEC-OVA to *Ly75*^+/+^ cells in the DZ at the 30, 46, and 60 h time points. This revealed a marked increase in representation of cells in S-phase cluster 2 as cells *Ly75*^+/+^ are positively selected in the LZ at 12 h (empty arrowheads in Fig. 2D). Enrichment in cluster 2 continued as cells transitioned to the DZ at 30 h, after which the dominant S-phase phenotype shifted to cluster 6 at 46 h and then cluster 3 at 60 h post DEC-OVA (filled arrowheads in Fig. 2D). Pairwise comparisons of the three S-phase clusters by gene set enrichment analysis (GSEA) using the “hallmark” signatures from the mSigDB database (Mootha et al., 2003; Subramanian et al., 2005) revealed expression of E2F and c-Myc target genes as the most consistent differences between the three clusters (Fig. S2A). Cluster 2, enriched in LZ-phenotype cells from the 12 and 30 h timepoints, showed high expression of *Myc* and of c-Myc- and mTORC1-dependent transcriptional signatures (Figs. 2E, and S2B) (Peng et al., 2002; Schuhmacher et al., 2001). Given the strong association between these signatures and positive selection (Calado et al., 2012; Dominguez-Sola et al., 2012; Ersching et al., 2017; Finkin et al., 2019; Luo et al., 2018; Victora et al., 2010), this pattern suggests that cluster 2 consists primarily of recently-selected B cells in “reactive” S phase, either on their way to, or soon after DZ reentry. Cells in clusters 3 and 6, on the other hand, showed noticeably reduced levels of *Myc* and c-Myc- and mTORC1-dependent gene signatures while maintaining expression of E2F target genes (Figs. 2E and S2B), as expected if they were entering the cell cycle by inertia while being physically segregated from mitogenic signals located in the LZ. Kinetic analysis of *Myc* and c-Myc and E2F target gene signatures confirmed that E2F target gene expression levels remained unaltered between 30 and 46 h post DEC-OVA, while mRNA expression and activity of Myc decreased (Figs. 2F-G and S2C). E2F signatures began to decay only at the 60 h time point, as inertial cycling defined by DZ S-phase entry (Fig. 1B,C) is already subsiding. Thus, unlike their LZ counterparts, DZ B cells appear to engage in a distinct mode of cell cycling that does not require continued expression of the *Myc* gene or its protein function to maintain E2F activity. This suggests that regulators of E2F that are down-stream of c-Myc and other selection-dependent signals may be required to sustain proliferation after positively-selected GC B cells transition to the DZ.

**Figure 2.**
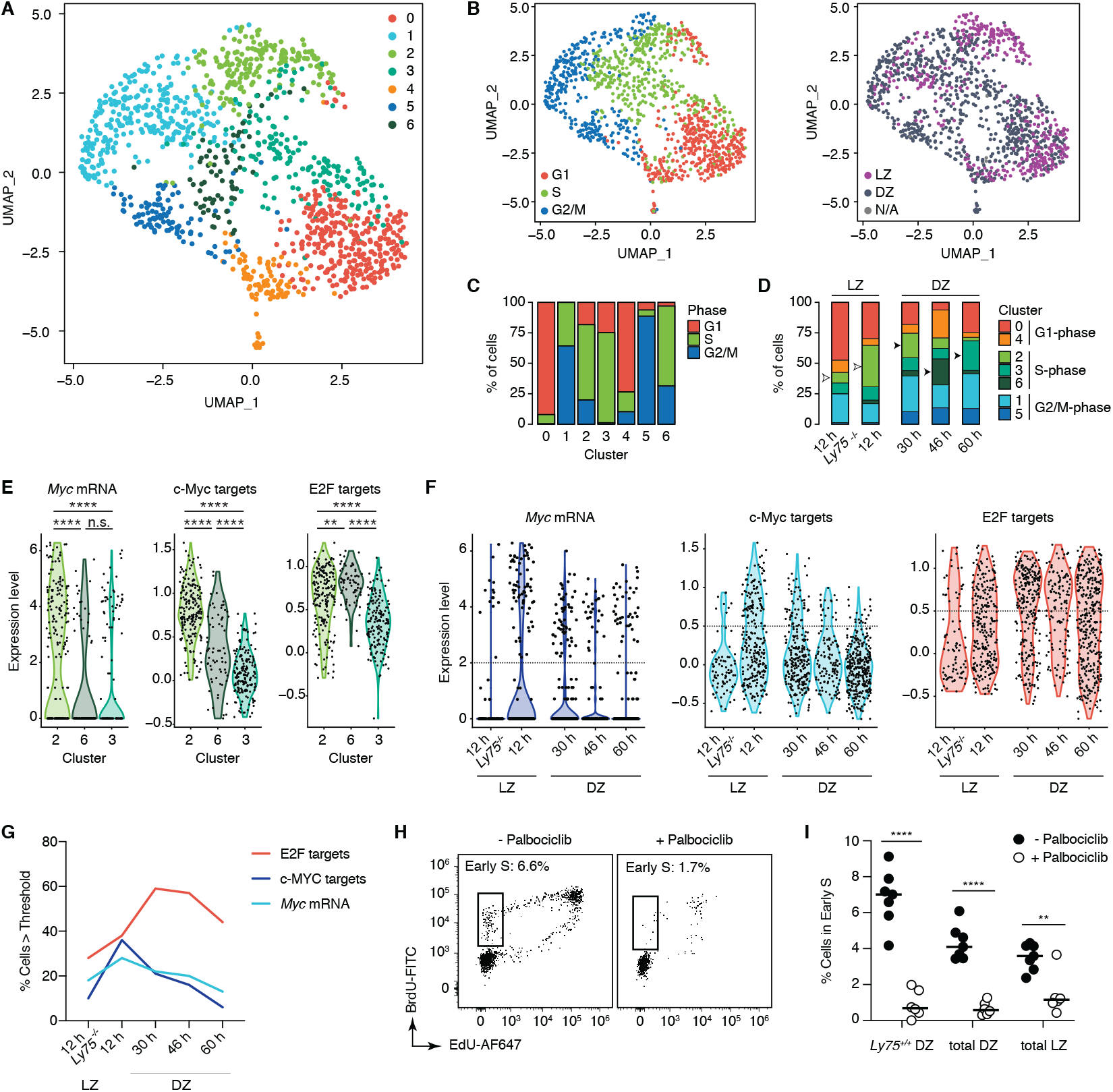
Single-cell transcriptomic analysis of GC B cells undergoing positive selection. (A) Uniform Manifold Approximation and Projection (UMAP) plot displaying 1220 cells colored by shared nearest neighbor (SNN) clusters collected (default Wilcox test; See Methods). Cells were collected from four independent experiments, see Table S1. (B) Distribution of cell cycle phase (left) and LZ/DZ phenotypes (right). (C) Distribution of cell cycle phase in clusters. (D) Changes in distribution of clusters grouped by cell cycle phase over the DEC-OVA-induced selection time course. (E-F) Expression of *Myc* mRNA, c-Myc and E2F target gene signatures in Clusters 2, 3, and 6 (E) or in DEC-OVA timepoints with indicated zonal phenotypes (F). Dotted line indicates the threshold used for quantification in (G). See Fig. S2C for a complete list of p values. (H) S phase entry of *Ly75*^+/+^ B1-8^hi^ cells in the DZ, 12 h post-treatment with Palbociclib, a CDK4/6 inhibitor, or vehicle (PBS). Quantification in (I) includes all LZ and DZ B1-8^hi^ cells, in addition to positively selected *Ly75*^+/+^ B1-8^hi^ cells in the DZ. **p < 0.01, ****p < 0.0001, nonparametric Mann-Whitney test, compared to the PBS-treated control group. Bars indicate median. Data pooled from two independent experiments.

A major regulator of E2F activity are the D-type cyclins, which in partnership with cyclin dependent kinases (CDKs) 4 and 6, activate E2F by phosphorylation of its negative regulator RB (Musgrove et al., 2011). To test whether D-type cyclins could be responsible for allowing the progression of inertial cell cycles, we treated mice with the inhibitor of CDK4/6 palbociclib 36 h after inducing positive selection of GC B cells with DEC-OVA and analyzed cell cycle progression 12 h later. Unlike blockade of MHC-II or CD40L (Fig. 1D-H), palbociclib treatment strongly inhibited S-phase entry in all GC B cells, including those undergoing inertial cycling in the DZ (Fig. 2H-I). Thus, inertial S-phase entry, while not dependent on Tfh-mediated signals, still requires activity of CDK4/6.

### Ccnd3 but not Ccnd2 is required for DZ inertial cycling

Since B cells express exclusively cyclins D2 and D3 (encoded by *Ccnd2* and *Ccnd3*, respectively) upon mitogenic stimulation (Reid and Snow, 1996; Solvason et al., 1996), we sought to determine the relative contribution of these two cyclins to the CDK4/6-dependency of inertial cycles. *Ccnd2* mRNA was detectable primarily in the subset of cells undergoing S phase in the LZ in the c-Mychigh cluster 2 (Fig. S3A). In agreement with our previous reports, *Ccnd2* was higher in *Ly75*^+/+^ cells at 12 h post-DEC-OVA as well as in the LZ in general (Dominguez-Sola et al., 2012; Victora et al., 2010) (Fig. S3B-C). To investigate the function of cyclin D2 in GC B cells, we generated *Ccnd2* knockout mice using CRISPR/Cas9-mediated genome editing in zygotes to introduce a 4-bp deletion/frameshift in exon 1 of the gene (Fig. S3D). *Ccnd2*^−/−^ mice lacked peritoneal B-1a cells (Fig. S3E-F) and females were unable to produce progeny (data not shown), as shown previously using an independently generated knockout strain (Sicinski et al., 1996; Solvason et al., 2000), confirming that ours is a null allele. We adoptively transferred a 1:1 mixture of *Ccnd2*^+/+^ and *Ccnd2*^−/−^ B1-8^hi^ cells into WT hosts, which we then immunized with NP-OVA to generate GCs (Fig. S3G) and found that zonal distribution was preserved in *Ccnd2*^−/−^ B1-8^hi^ B cells (Fig. S3H-I). However, rather than reducing the ability of GC B cells to cycle, loss of cyclin D2 led to a small but consistent increase in the proportion of B1-8^hi^ cells entering S phase in both LZ and DZ (Fig. S3J-L). Despite this increase, absence of cyclin D2 showed no clear effect on the competitiveness of GC B cells over time when compared to *Ccnd2*-sufficient B cells within the same GC (Fig. S3M-N). These experiments suggest that, although cyclin D2 can have a negative effect on the ability of GC B cells to enter cell cycle, it is not required for inertial cycling in the DZ.

In contrast to the dynamic behavior of *Ccnd2*, our scRNA-seq dataset showed that *Ccnd3* mRNA amounts were stably high throughout the LZ/DZ cycle and over our time course of DEC-OVA-induced selection, with only slight increases at the 30 h time point and in the DZ in general (Fig. 3A-C). Despite these modest changes in mRNA expression, cyclin D3 protein levels were substantially higher in the DZ (Fig. 3D), in agreement with previous reports based on histology (Peled et al., 2010). Also consistently with prior reports (Cato et al., 2011; Peled et al., 2010), we found that loss of *Ccnd3* led to dramatically reduced GC B cell frequency (Fig. 3E-F). This is unlikely to be due to a general defect in proliferation, given that *Ccnd3*^−/−^ B cells proliferate normally in vitro and at pre-GC stages in vivo (Cato et al., 2011; Peled et al., 2010). Closer examination of *Ccnd3*^−/−^ GCs showed that decreased GC size was primarily caused by a marked reduction in the proportion of B cells with a DZ phenotype (Fig. 3G-H), suggesting a strong block in the ability of *Ccnd3*^−/−^ B cells to undergo inertial cycling. This phenotype was identical when GCs were generated by adoptive transfer of *Ccnd3*^−/−^ B1-8^hi^ cells into WT hosts (Fig. 3I),resulting in a significantly decreased DZ/LZ ratio (Fig. 3J-K) and an almost complete loss in S phase initiation in the DZ with only a minor decrease of fitness in the LZ (Fig. 3J, L-N). The requirement for cyclin D3 is therefore GC B cell-intrinsic and largely restricted to the DZ. *Ccnd3*^−/−^ B1-8^hi^ cells in the DZ also showed increased forward scatter, a surrogate measure of cell size (Fig. 3O-P). Since cell size increases prior to the initiation of inertial cycles (Ersching et al., 2017; Finkin et al., 2019), this finding suggests that the inability of *Ccnd3*^−/−^ B cells to undergo inertial cycling results in their failure to lose cellular mass by division. We conclude that cyclin D3 is cell-intrinsically required for GC B cells to enter S phase specifically in the DZ, while being dispensable for S phase entry immediately down-stream of T cell-mediated selection in the LZ. This pattern implicates cyclin D3 as a nonredundant mediator of inertial cell cycles.

**Figure 3.**
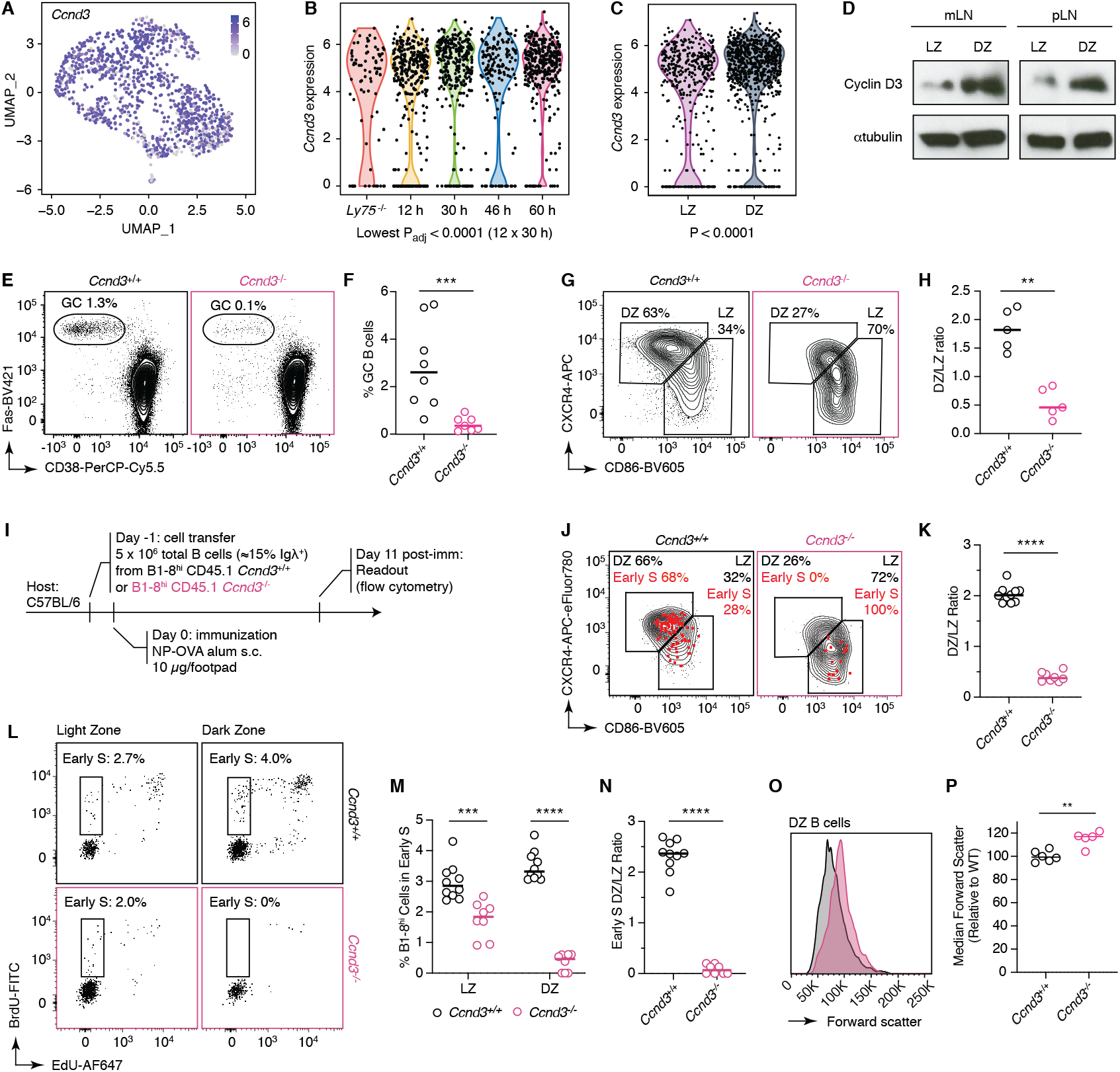
Inertial B cell cycling requires cyclin D3. (A) Expression of *Ccnd3* in UMAP dimension (A), over time after DEC-OVA immunization (B), and in LZ or DZ (C). P-values in (B) are for Kruskall Wallis test with Dunn’s multiple comparisons test. Other significant P values are: < 0.001 (12 x 60h); 0.013 (*Ly75*^−/−^ vs. 30 h), and 0.015 (*Ly75*^−/−^ vs. 60 h). (D) Immunoblots of whole cell lysates of LZ or DZ cells sorted from popliteal or mesenteric lymph nodes (pLN or mLN, respectively). (E-H) Staining for GC (E) and DZ/LZ (G) in WT (*Ccnd3*^+/+^) or cyclin D3 mutant (*Ccnd3*^−/−^) mice that were immunized s.c. in the hind footpad for pLN, quantified in (F). (I) Experimental setup for induction of GCs containing WT (*Ccnd3*^+/+^) or cyclin D3 mutant (*Ccnd3*^−/−^) B1-8^hi^ cells. (J-P) DZ and LZ staining of all GC B cells (black) or cells entering S phase (red) (J), S phase entry (L), forward scatter (O) of WT (*Ccnd3*^+/+^) or cyclin D3 mutant (*Ccnd3*^−/−^) B1-8^hi^ cells, quantified in (K), (M), (N), and (P). *p < 0.05, **p < 0.01, ***p < 0.001, ****p < 0.0001, n.s. non-significant, nonparametric Mann-Whitney test. Bars indicate median. Data pooled from three (E-H) or two (I-P) independent experiments.

### Cyclin D3-mediated DZ inertial cycling links GC positive selection with clonal expansion

To determine the relative importance of reactive LZ cycles vs. inertial cell DZ cycles for clonal expansion, we asked whether cell proliferation defects seen in *Ccnd3*^−/−^ DZ B cells could be rescued by forcing strong interaction with Tfh cells using DEC-OVA (Fig. 4A). Whereas *Ccnd3*^+/+^ *Ly75*^+/+^ B1-8^hi^ cells proliferated sufficiently to outnumber *Ly75*^−/−^ cells by 5-fold over a 60 h period, clonal expansion was much less efficient when *Ly75*^+/+^ B1-8^hi^ cells lacked cyclin D3. With the exception of one outlier mouse, the ratio of *Ly75*^+/+^ to *Ly75*^−/−^ cells increased only slightly, if at all, over input levels (Fig. 4B-C). Failure of *Ccnd3*^−/−^ B cells to clonally expand was also accompanied by failure to accumulate in the DZ (Fig. 4D-E), although histology showed that *Ccnd3*^−/−^ cells were able to at least access the DZ anatomically at 36 h post-DEC-OVA (Fig. 4F). Even so, the proportion of early S-phase cells among the *Ccnd3*^−/−^ population dropped precipitously upon transition from LZ to DZ, as expected given the inability of cyclin D3-deficient B cells to sustain proliferation by inertia (Fig. 4G-K). Thus, the residual expansion of *Ccnd3*^−/−^ *Ly75*^+/+^ B1-8^hi^ seen in some mice can be attributed primarily to reactive cell cycling taking place in the LZ, in response to continued signals delivered by Tfh cells. We conclude that strong signaling from Tfh cells in response to DEC-OVA cannot overcome the requirement for cyclin D3 in driving DZ proliferation, and that, in the absence of inertial cycling, reactive cell cycles triggered directly by Tfh-derived signals are not sufficient to sustain large proliferative bursts.

**Figure 4.**
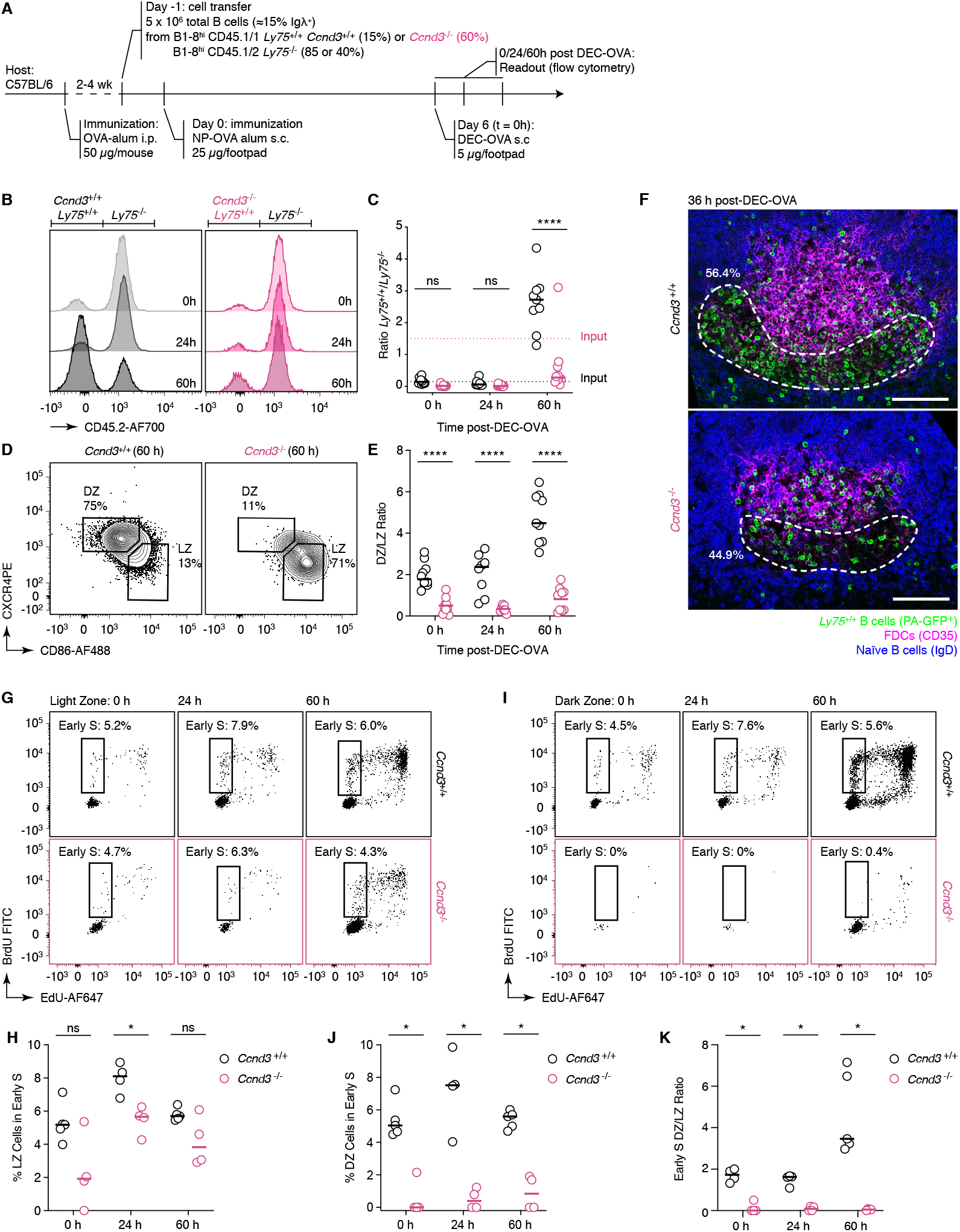
Increased Tfh cell help cannot compensate for loss of cyclin D3. (A) Experimental setup for DEC-OVA-induced positive selection of *Ly75*^+/+^ cells that are either WT (*Ccnd3*^+/+^) or cyclin D3 mutant (*Ccnd3*^−/−^). Note that due to of lack of competitiveness in early GC stages, *Ccnd3*^−/−^ B1-8^hi^ cells have to be transferred at a higher proportion than *Ccnd3*^+/+^ B1-8^hi^ cells. (B-E) Clonal expansion (B) and DZ/LZ staining (D) of WT (*Ccnd3*^+/+^) or cyclin D3 mutant (*Ccnd3*^−/−^) *Ly75*^+/+^ cells over time after DEC-OVA immunization, quantified in (C) and (E). (F) Immunofluorescence showing the migration of WT (*Ccnd3*^+/+^) or cyclin D3 mutant (*Ccnd3*^−/−^) *Ly75*^+/+^ cells 36 h post DEC-OVA-immunization in pLN GCs. Dotted area indicates the DZ marked by the absence of IgD (Naïve B cell marker) and CD35 (FDC marker). Percentage of *Ly75*^+/+^ cells within the DZ out of the entire GC is quantified. (G-K) S phase entry of positively selected *Ly75*^+/+^ B1-8^hi^ cells that are either WT (*Ccnd3*^+/+^) or cyclin D3 mutant (*Ccnd3*^−/−^) in the LZ (G) or DZ (H), quantified in (I), (J), and (K). *p < 0.05, **p < 0.01, ***p < 0.001, ****p < 0.0001, n.s. non-significant, nonparametric Mann-Whitney test, compared to WT (*Ccnd3*^+/+^). Bars indicate median. Data pooled from four independent experiments.

### Cyclin D3 controls DZ inertial cycling in a dose-dependent manner

A potential mechanism for how the strength of the initial T cell-B cell interactions in the LZ determines the number of rounds of division that selected B cells undergo in the DZ (Gitlin et al., 2014), is that DZ B cells translate the memory of their interactions with Tfh cells into protein amounts of a cell cycle regulator. If cyclin D3 follows such a pattern, it would be predicted that GC B cells with a higher capacity to produce cyclin D3 protein would be at a competitive advantage due to increased proliferation in the DZ. To test this, we directly competed B1-8^hi^ B cells carrying either one or two intact alleles of *Ccnd3* by adoptive transfer of a 1:1 mixture of *Ccnd3*^+/−^ and *Ccnd3*^+/+^ B1-8^hi^ B cells into the same recipient mice, which were then immunized with NP-OVA in alum and assayed for relative abundance of these two populations (Fig. 5A). Whereas both *Ccnd3*^+/+^ and *Ccnd3*^+/−^ B1-8^hi^ cells were found at a similar ratio in early GCs at day 7 post-immunization, the proportion of *Ccnd3*^+/−^ B1-8^hi^ cells decreased gradually over time, such that these cells were completely eliminated in 3/7 mice by day 14 post-immunization (Fig. 5B-C). Thus, the reduction in cyclin D3 dosage associated with heterozygosis is sufficient to impose a gradual but clear loss of this population form the GC. Lack of competitiveness of heterozygous B cells was associated with a slight reduction in the DZ/LZ ratio, already observable at day 7 post-immunization (Fig. 5D-E), and a decrease in the proportion of cells entering S phase in the DZ, but not in the LZ (Fig. 5F-H). To extrapolate the loss of inertial proliferative capacity among *Ccnd3*^+/−^ GC B cells from our direct competition data, we simulated this experiment in silico using a previously published agent-based model of GC selection that includes T cell control over the number of cell cycles carried a B cell undergoes upon positive selection (Meyer-Hermann, 2020; Meyer-Hermann et al., 2012). We modeled loss of *Ccnd3* expression as a reduction in the maximum number of divisions a *Ccnd3*^+/−^ GC B cell can complete upon positive selection. We varied the relationship between the degree of T cell help (modeled as intensity of c-Myc activation) and the number of divisions as illustrated by the curves shown in Figure 5I. The experimentally measured kinetics of *Ccnd3*^+/−^ GC B cells were best reproduced when the maximum number of divisions for *Ccnd3*^+/−^ GC B cells in silico was reduced to 72% of WT, with a residual sum of squares (RSS) of 0.14 (Fig. 5J). In silico, a reduction in responsiveness of this magnitude in *Ccnd3*^+/−^ GC B cells was accompanied by a reduction in the number of cell cycles per cell to approximately 84% of WT levels at day 8 post-immunization, which is compatible with our EdU/BrdU incorporation data (Fig. 5K). We conclude that cyclin D3 dose-dependently controls the number of inertial cycles a B cell will undergo in the DZ. The sensitivity of GC B cells to loss of even a single allele of *Ccnd3* suggests that cyclin D3 protein abundance may serve as a molecular bridge linking the cumulative signal a B cell receives from T cells in the LZ and the number of cycles this cell can execute in the DZ.

**Figure 5.**
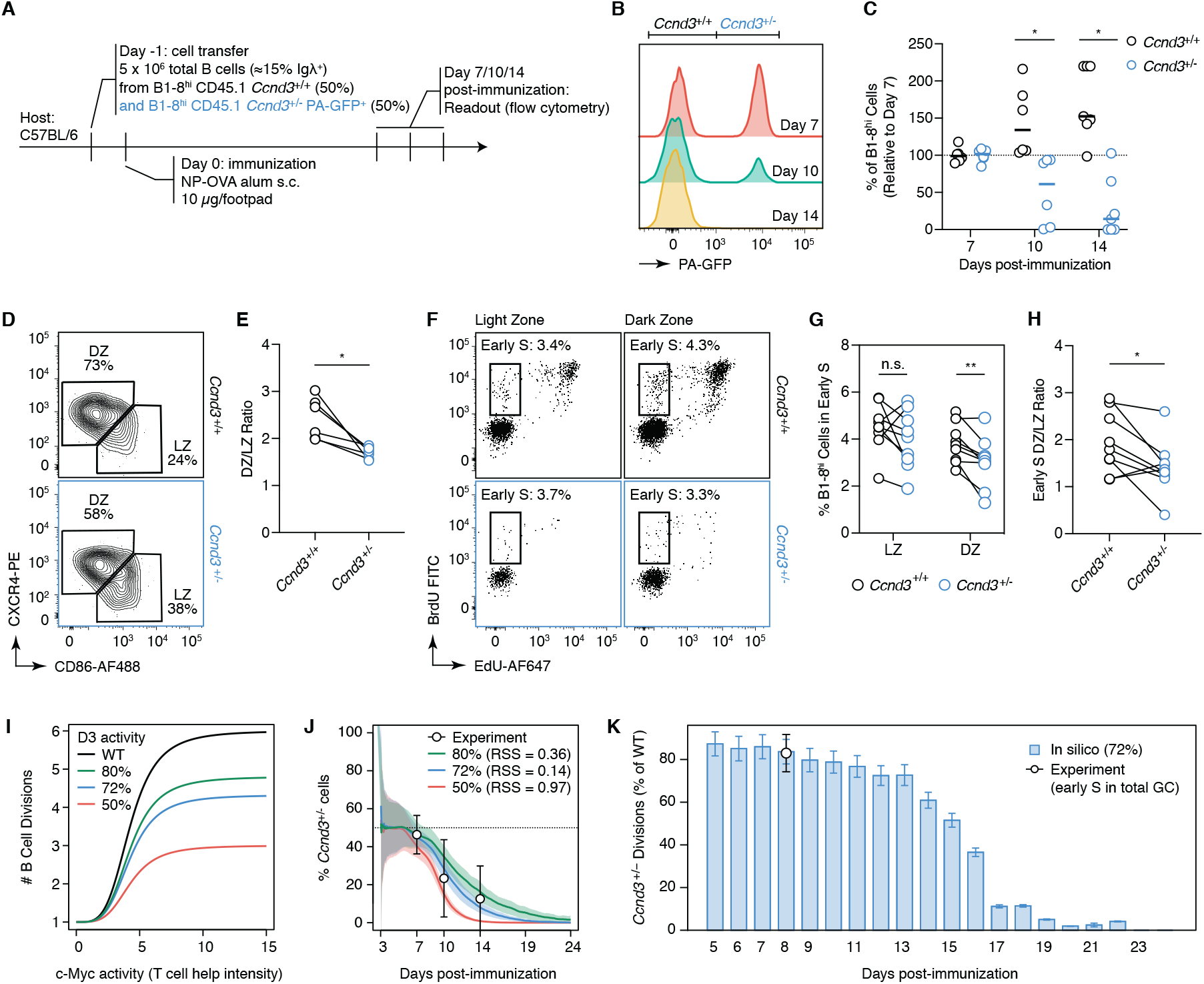
Cyclin D3 controls inertial cell cycling in a dose-dependent manner. (A) Experimental setup for induction of GCs containing mixtures of B1-8^hi^ cells with a full (*Ccnd3*^+/+^) or reduced (*Ccnd3*^+/−^) dose of cyclin D3. (B) Clonal expansion of B1-8^hi^ cells with a full (*Ccnd3*^+/+^) or reduced (*Ccnd3*^+/−^) dose of cyclin D3 over time, quantified in (C) relative to Day 7. (D-H) DZ and LZ staining in GC B1-8^hi^ cells 7 days after NP-OVA immunization, quantified in (E). Dotted line indicates averaged DZ/LZ ratio of total GC B cells. (F) S phase entry of B1-8^hi^ cells with a full (*Ccnd3*^+/+^) or reduced (*Ccnd3*^+/−^) dose of cyclin D3 in Light or Dark zone 8 days after NP-OVA immunization, quantified in (G) and (H). (I) Relationship between strength of T cell help (modeled as c-Myc signal intensity) and number of B cell divisions in the different models used to estimate loss of function in *Ccnd3*^+/−^ GC B cells. The maximum number of divisions allowed to *Ccnd3*^+/−^ cells corresponds to 50% (red) or 72% (blue) or 80% (green) of the maximum number of divisions of the WT. (J) *Ccnd3*^+/−^ kinetics in silico. Loss of *Ccnd3*^+/−^ cells as a fraction of the GC population over time in silico. The maximum number of divisions of *Ccnd3*^+/−^ GC B cells was fixed to 50% (red line), 72% (blue line), or 80% (green line) of the WT value. Mean (solid lines) and standard deviation (shaded area) over 60 GC simulations. Experimental mean ± SD is shown as white circles. RSS, residual sum of squares. (K) *Ccnd3*^+/−^ divisions in silico. The mean number of divisions of *Ccnd3*^+/−^ cells is shown as a percentage of WT at the indicated days. Reported data result from simulations with the maximum number of divisions of the *Ccnd3*^+/−^ GC B cells fixed to 72% of the WT maximum number of divisions. Mean and standard deviation over 60 GC simulations. White circle represents experimental data. *p < 0.05, **p < 0.01, paired t test. Bars indicate median. Data pooled from at two independent experiments.

### A lymphoma-associated mutation that stabilizes cyclin D3 promotes DZ inertial cycling and drives clonal B cell lymphoproliferation

A parallel line of evidence pointing to the importance of cyclin D3 in GC proliferation is the presence of a series of *Ccnd3* mutations that stabilize cyclin D3 protein in roughly 40% of cases of sporadic Burkitt lymphoma (Casanovas et al., 2004; Schmitz et al., 2014; Schmitz et al., 2012; Sonoki et al., 2001). Because Burkitt lymphoma cells closely resemble DZ B cells in gene expression (Caron et al., 2009; Victora and Mouquet, 2018), we hypothesized that stabilization of cyclin D3 by means of a lymphoma-associated mutation may increase the propensity of GC B cells to undergo inertial cycles in the DZ. To test this, we used CRISPR/Cas9-mediated genome editing to generate a *Ccnd3* allele encoding a version of cyclin D3 protein that is stabilized by the replacement of a phosphorylatable threonine (T283A) in its C-terminal domain (Fig. 6A). This mutation prevents phosphorylation of T283, which would otherwise promote nuclear export and proteasomal degradation of cyclin D3 (Casanovas et al., 2004; Cato et al., 2011). When introduced into the genome, the T283A mutation alone did not cause any overt anomaly in B celldevelopment, with the possible exception of a slight increase in the percentage of pre/pro B cells in the bone marrow, nor did it lead to spontaneous GC formation in the spleens of young adult mice (Fig. S4A,D). Nonetheless, we found that, upon immunization with NP-OVA, *Ccnd3*^*T283A*/+^ mice generated GCs that were approximately twice as large as WT GCs and had a markedly higher DZ/LZ ratio, suggestive of increased DZ proliferation (Fig. 6B-E). Accordingly, flow cytometric measurement of EdU/BrdU incorporation showed a ~50% increase in GC B cells entering S phase in the DZ, but no increase in S phase entry in the LZ (Fig. 6F-H). To confirm that the proliferative effect of cyclin D3 on the GC is B cell-intrinsic, we generated mixed chimeras in which bone marrow cells from either WT or *Ccnd3*^*T283A*/+^ mice were mixed with those from B cell-deficient JHT mice at a 20:80 ratio.Consistent with the analysis of *Ccnd3^T283A/+^* mice, chimeras with *Ccnd3*^*T283A*/+^ B cells showed increased GC size, DZ expansion, and S-phase entry in the DZ, but not in the LZ, upon immunization with NP-OVA in alum (Fig. 6C,E,G,H). This phenotype was recapitulated upon immunization with keyhole limpet hemocyanin (KLH), ruling out any potential effects on of the hypomorphic immunoglobulin λ light chain of the SJL strain in which *Ccnd3*^T283A^ mice were originally generated on the normally Igλ-dominated response to NP-OVA (Fig. S4E-G). Moreover, *Ccnd3*^*T283A*/+^ DZ B cells showed a decrease in size as measured by forward scatter, mirroring the phenotype found in *Ccnd3*^−/−^ B cells (Fig. 6I-J). This indicates that the increased proliferation of *Ccnd3*^*T283A*/+^ B cells in the DZ extends to the extreme of what these cells are metabolically programmed to support (Ersching et al., 2017). Accordingly, adoptive transfer of a 1:1 mixture of *Ccnd3*^+/+^ and *Ccnd3*^*T283A*/+^ B1-8^hi^ B cells into the same recipient mice followed by immunization with NP-OVA in alum showed that the representation of *Ccnd3*^*T283A*/+^ B1-8^hi^ cells in GCs increased gradually but only slightly over time, suggesting that the proliferative capacity endowed by the *Ccnd3*^*T283A*^ mutation remains limited by metabolic capacity (Fig. 6K-M). Together, these data indicate that T283A mutation in cyclin D3 specifically promotes inertial DZ cell cycle progression in a B cell-intrinsic manner, leading to increased competitiveness of mutant B cells in the GC.

**Figure 6.**
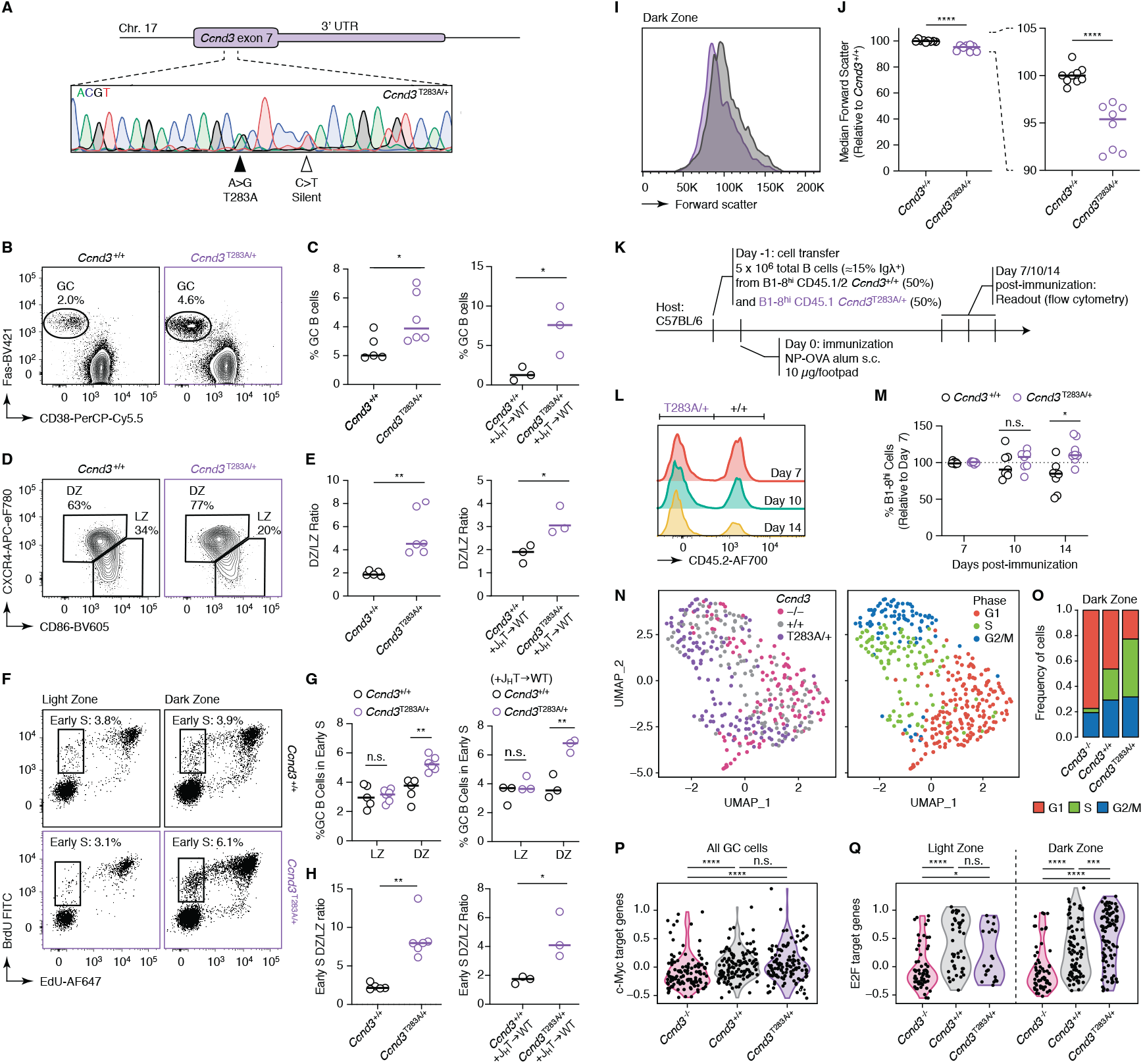
A lymphoma-derived mutation in *Ccnd3* exclusively promotes inertial cell cycle entry. CRISPR/Cas9-mediated gene targeting strategy to introduce a single nucleotide substitution resulting in a T283 to A mutation and a silent mutation to remove the PAM motif to prevent Cas9 recutting. (C-H) (left) Staining for GC (B), DZ/LZ (D), and S phase entry (F) of GC B cells in pLNs from WT (*Ccnd3*^+/+^) or *Ccnd3*^*T283A*/+^ mice, quantification in (C, E, G, H): (left) WT (*Ccnd3*^+/+^) or *Ccnd3*^*T283A*/+^ mice or (right) BMCs generated with BM cells isolated from WT (*Ccnd3*^+/+^) or *Ccnd3*^*T283A*/+^ and B cell-deficient JHT mice at a 20:80 ratio. (I) Forward scatter of DZ B cells sorted from pLNs of WT (*Ccnd3*^+/+^) or *Ccnd3*^*T283A*/+^ mice, quantification of median forward scatter of DZ B cells in (J). (K) Experimental setup for induction of GCs containing mixtures of B1-8^hi^ cells with a full (*Ccnd3*^+/+^) or increased (*Ccnd3*^*T283A*/+^) dose of cyclin D3. (L) Clonal expansion of B1-8^hi^ cells with a full (*Ccnd3*^+/+^) or reduced (*Ccnd3*^*T283A*/+^) dose of cyclin D3 over time, quantified in (M) relative to Day 7. (N) UMAP plot colored by genotype (left) or cell cycle phase (right). (O) Distribution of cell cycle phase in each genotype. (P-Q) Quantification of c-Myc (P) or E2F (Q) target gene signatures in in each genotype. *p < 0.05, **p < 0.01, ***p < 0.001, n.s. non-significant, nonparametric Mann-Whitney test, compared to WT (*Ccnd3*^+/+^) (C, E, G, H, J, P, Q), or paired t test comparison between WT (*Ccnd3*^+/+^) or *Ccnd3*^*T283A*/+^ cells in the same animal (M). Bars indicate median. Data pooled from two (B-H), three (I-J), two (L-M), or one (N-Q) independent experiments.

To understand the effect of cyclin D3 dosage on GC B cell cycles at the transcriptional level, we performed scRNAseq on individually sorted *Ccnd3*^+/+^, *Ccnd3*^−/−^, and *Ccnd3*^*T283A*/+^ GC B cells. Whereas these cells segregated primarily by cell cycle phase rather than genotype (Fig. 6N), we found that an increase in cyclin D3 dosage resulted in a marked shift in the fraction of DZ B cells from G1 to S and G2/M phases (Fig. 6O), suggesting that DZ B cells with high expression of cyclin D3 are poised to undergo inertial cycling and swiftly transit between S and M phases with only a brief or indiscernible G1 phase. Consistent with the inertial cell cycling phenotype revealed by DEC-OVA treatment, E2F target gene expression levels in single cells increased in a cyclin D3dose-dependent manner in the DZ, despite comparable expression of c-Myc target genes across all genotypes (Fig. 6P-Q). We conclude that, whereas c-Myc can behave as a “charger” that prompts reactive cycling in the LZ, sustained E2F target gene expression downstream of cyclin D3 dosage provides the mechanism enabling inertial cycling in the DZ.

The finding that a common lymphoma mutation increases the ability of GC B cells to undergo inertial cell cycles (Schmitz et al., 2012) suggests that lymphomas may specifically coopt the mito-gen-independent proliferative program that characterizes the GC DZ to achieve sustained and rapid proliferation. To determine the long-term effects of the *Ccnd3*^T283A^ mutation on B cell proliferation and lymphomagenesis, we aged cohorts of *Ccnd3*^*T283A*/+^ mice for up to ~1.5 years. The T283A mutation in cyclin D3 was not sufficient to cause overt morbidity or early mortality (Fig. 7A). However, we found that B cells from *Ccnd3*^*T283A*/+^ mice, but not their WT littermate controls, developed a population of Fas+ CD38+ double-positive (DP) B cells whose frequency increased over time (Fig. 7B-C). Sequencing of the *Igh* genes of DP B cells revealed that this population became largely monoclonal as its size increased (Fig. 7D-E), implicating *Ccnd3*^T283A^ as a driver of B cell lymphoproliferation, but also suggesting that additional mutations are likely required to achieve this phenotype. In the two mice with the largest DP population, expanded clones were class-switched, heavily mutated, and were also present within populations with a more typical GC phenotype (Fas+ CD38-), indicative of GC origin (Fig. 7D, F). In the most extreme case, DP and GC B cells isolated from various tissue sources together comprised a phylogenetic tree that was both heavily expanded and highly branched, indicating continual proliferation and somatic hypermutation (Fig. 7F). In addition to the clonal DP lymphoproliferative phenotype, aged *Ccnd3*^T283A^ mice displayed other key signs of hematopoietic malignancy that were not observed in any of their WT counterparts. These included expansion of abnormal lymphocyte populations, enlarged secondary lymphoid organs with disrupted morphology, and lymphocytic infiltration into non-lymphoid organs, such as the liver, lungs, and kidneys (Fig. S5). While the variable nature of the malignant phenotypes observed precluded us from carrying out a systematic characterization of these outgrowths, these data confirm the oncogenic potential of *Ccnd3*^T283A^ in lymphoid cells. We conclude that the hyperactivation of cyclin D3 by the T283A mutation promotes the acquisition of a clonal lymphoproliferative phenotype by B cells, and more generally favors malignant transformation of lymphocyte populations.

**Figure 7.**
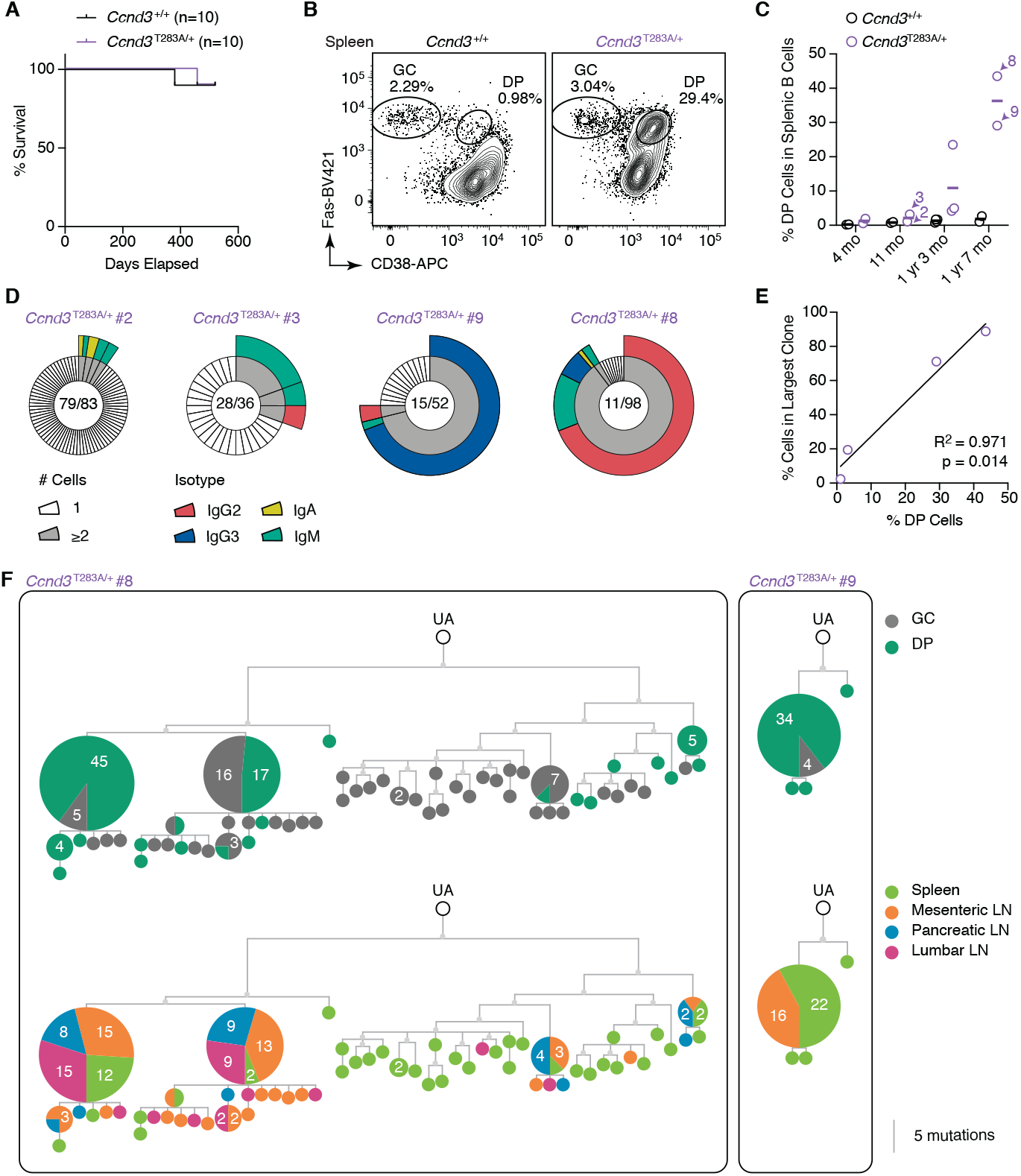
A lymphoma-derived mutation in *Ccnd3 drives clonal B cell lymphoproliferation*. (A) Survival curve of WT (*Ccnd3*^+/+^) or *Ccnd3*^*T283A*/+^ mice. (B) Staining for GC/DP in splenic B cells, quantified in (C) by the age of mice. Mice indicated with arrow-heads/numbers were chosen for sequencing of the *Igh* genes. (D) Pie chart showing clonal distribution of DP cells from spleen and mesenteric LNs. *Ccnd3*^*T283A*/+^ #8 also includes DP cells from pancreatic and lumbar LNs. Each slice in inner rings represents one clone indicated by a distinct V(D)J rearrangement; outer rings are isotypes represented in each clone found in at least two cells. Numbers are (clones observed)/(cells sequenced). (E) Proportional relationship between % DP cells among B220+ B cells and % cells in the largest clone identified in each mouse. (F) Trees showing phylogenetic relationships between VH sequences of cells from selected clones and their unmutated VH ancestor (UA). Numbers inside cells indicate frequency of a particular sequence observed in combination with a GC/DP phenotype (top) or across various tissues (bottom) from the same mouse. Scale bar indicates 5 mutations.

## Discussion

A central question in GC biology is how antigen-dependent signals delivered to GC B cells lead to the rapid clonal expansion that characterizes positive selection of higher-affinity SHM variants. Within the framework of Tfh cell-driven B cell selection, this question can be translated into how Tfh cell help delivered in the LZ controls the number of cell cycles a B cell subsequently undergoes after transition to the DZ (Mesin et al., 2016). Because this number is proportional to the “push” a B cell receives from Tfh cells while in the LZ (Gitlin et al., 2015; Gitlin et al., 2014; Meyer-Hermann et al., 2012; Victora et al., 2010), and our antibody blocking experiments formally rule out that DZ-initiated cell cycles depend on continued presence of Tfh cell help, we refer to DZ cell cycles as “inertial.” According to such a model, GC B cells would alternate between “reactive” cell cycles, in which a single S phase entry occurs in the LZ in direct response to mitogenic signals delivered by Tfh cells, and inertial cycles, whereby B cells reenter S phase in the DZ a number of times proportional to the Tfh cell help they received in the LZ, but independently of continued Tfh cell help.

Our findings show that reactive and inertial cell cycles differ with respect to their molecular requirements. A number of studies have pointed towards c-Myc as a critical regulator of the DZ proliferative burst (Calado et al., 2012; Dominguez-Sola et al., 2012; Ersching et al., 2017; Finkin et al., 2019). Previous work suggests that c-Myc partners with mTORC1 to fuel the cellular growth that precedes the proliferative burst and dose-dependently controls burst magnitude (Ersching et al., 2017; Finkin et al., 2019). However, c-Myc itself is not a driver of the DZ cell cycle, since its expression is predominantly found in the LZ (Calado et al., 2012; Dominguez-Sola et al., 2012; Victora et al., 2010). Our scRNA-seq experiments show that the expression of *Myc* and c-Myc-dependent genes in the DZ are largely restricted to cells in S phase at the earliest timepoint after strong positive selection in the LZ, whereas inertial cycling continues well beyond the disappearance of c-Myc-associated gene signature expression and is likely to involve signals that are still active during G2/M (Spencer et al., 2013). Thus, even though c-Myc is dose-dependently required for inertial cycling (Finkin et al., 2019), it is likely that c-Myc itself is analogous to a “charger” of the pre-division phenotype rather than to a “timer” (Bannard et al., 2013) or “counter” (Gitlin et al., 2014) of post-signal DZ divisions. Instead, inertial cycling seems to be fueled by the maintenance of E2F activity. Previous studies have reported stabilized expression of E2F transcription factors during mitosis in association with shortening of G1 and faster S phase entry in the subsequent cell cycle (Clijsters et al., 2019; Sigl et al., 2009). Thus, we propose that these molecular signatures may characterize DZ B cells that are fated to undergo one or more subsequent rounds of cell cycling by inertia. On the other hand, DZ B with low levels of E2F would be expected to pause in G1 and migrate back to the LZ for another round of antigen-driven selection.

We find that B cell-intrinsic expression of cyclin D3 is specifically required for inertial DZ cycles, whereas it plays at best a partly redundant role in the reactive cell cycles initiated in the LZ. Our experiments with a series of *Ccnd3* alleles with varying degrees of mRNA and protein expression show that cyclin D3 controls the extent of DZ B cell proliferation and the size of the GC DZ in a dose-dependent manner, with fully deficient cells being almost completely unable to initiate S phase in the DZ and heterozygous B cells showing roughly 70% of the proliferative ability of WT cells. This suggests an instructive role for cyclin D3 in controlling GC B cell burst size. A possible mechanism for this control is that the intensity of a B cell’s interaction with Tfh cells is converted to a proportionate amount of cyclin D3 protein by increased translation of mostly unchanging levels of *Ccnd3* mRNA or by stabilization of cyclin D3 protein by preventing its degradation by the ubiquitin-proteasome system, potentially with an additional contribution of increased transcription of the *Ccnd3* gene. Previous work has shown that cyclin D1 levels carry through mitosis to accelerate subsequent S-phase entry (Min et al., 2020). Our data suggest that a similar mechanism, driven by cyclin D3, may be at play in DZ inertial cycles.

A clear indication of the importance of cyclin D3 to GC B cell proliferation, especially in the DZ, is the frequent presence of *Ccnd3*^T283A^ or functionally equivalent mutations in Burkitt lymphoma (Schmitz et al., 2014; Schmitz et al., 2012), the most DZ-like of B cell malignancies (Caron et al., 2009; Victora et al., 2012). The strong selection for this mutation in Burkitt lymphoma suggests that these cells may take advantage of the GC inertial cycling program to amplify the number of divisions they can undergo in response to the *Myc*-*Igh* translocation that defines this malignancy. Whereas mice bearing the *Ccnd3*^T283A^ mutation did not succumb to B cell lymphoma, they did develop large age-related expansions of B cell clones that spanned the typical GC (FAS+ CD38-) and an unusual FAS+ CD38+ DP phenotype, with isotype and somatic mutation patterns suggestive of GC origin. Similar DP B cells comprised 4 of 9 CD19+ lymphomas arising in mice with p53-deficient B cells upon *Plasmodium* infection (Robbiani et al., 2015). The finding that most cells in mice with large DP expansions derived from a single clone suggests that increased inertial proliferation downstream of the cyclin D3 T283A mutation may predispose B cells to acquire further somatic mutations or chromosomal rearrangements that lead to the lymphoproliferative phenotype, and may also more rapid proliferation downstream of the acquisition of secondary mutations. The oncogenic potential of this mutation is underscored by the development of various forms of hematopoietic abnormalities in *Ccnd3*^T283A^ mice. Understanding the molecular drivers of inertial vs. signal-dependent cell cycling and how and why these are coopted by lymphoid malignancies may shed light on vulnerabilities that are specific to inertially cycling cells, providing an avenue that could be exploited therapeutically.

Finally, from the perspective of GC selection, the failure of *Ccnd3*^−/−^ GC B cells to clonally expand even upon strong induction of selective signals by DEC-OVA suggests that combining reactive and inertial cell cycles is required for the “clonal busting” that leads to optimal expansion of higher-affinity B cell clones in physiological settings (Tas et al., 2016). GC B cells appear to balance clonal bursting with other, less exclusive forms of selection as a strategy to evolve high-affinity clones while maintaining some degree of clonal diversity (Amitai et al., 2017; Bannard and Cyster, 2017; Mesin et al., 2016; Tas et al., 2016). Control of inertial selection by cyclin D3 may play an important role in fine-tuning this balance.

## Author Contributions

All in vivo experiments were designed and performed by J.P., J.E., A.E., and G.D.V. T.B.R.C. carried out the bioinformatic analysis of scRNAseq data. L.M., S.J.A., J.O.-M., contributed additional scRNAseq libraries, under supervision of A.K.S. M.S. and M.M.-H. performed mathematical modeling. C.M. and A.M. contributed to the description of the malignant phenotype of *Ccnd3*^T283A^ mice, including histological analysis. J.P., T.B.R.C., and G.D.V. wrote the manuscript with input from all authors. G.D.V. supervised the work.

## Acknowledgements

We thank P. Sicinski (Dana-Farber Cancer Institute) and H. Lodish (Whitehead Institute) for *Ccnd3*^−/−^ mice, Kristie Gordon and Kalsang Chhosphel for FACS sorting, S. Markoulaki and R. Jaenisch (Whitehead Institute) and R. Norinsky (Rockefeller University) for CRISPR/Cas9 mouse generation, the Laboratory of Comparative Pathology (MSKCC) for mouse tissue processing and staining, and David Dominguez-Sola (Mount Sinai School of Medicine) for critical reading of our manuscript. This work was funded by NIH/NIAID grants R01AI139117 and R01AI119006, to G.D.V. and Starr Consortium Grant I11-0027 to A.M. and G.D.V. with additional support from NIH grant DP1AI144248 (Pioneer award) to G.D.V. J. P. is the Berger Foundation Fellow of the Damon Runyon Cancer Research Foundation (DRG-2353-19). J.E. is a Cancer Research Institute-Irvington postdoctoral fellow. M.S. was supported by the European Union’s Horizon 2020 research and innovation program under the Marie Skłodowska-Curie grant agreement no. 765158. J.O-M was supported by the Damon Runyon Cancer Research Foundation (DRG-2274-16) and Richard and Susan Smith Family Foundation. C.M. was supported by the LRF (Lymphoma Research Foundation) Post-doctoral Fellowship, the LLS (The Leukemia & Lymphoma Society) Special Fellow Award and the ASH Research Restart Award for early career investigators in hematology. A.M. was supported by NCI grant R35CA220499. A.E. is a Ramon y Cajal Awardee MICIU/AEI (RYC-2013-13546). A.K.S. was supported by the Searle Scholars Program, the Beckman Young Investigator Program, a Sloan Fellowship in Chemistry, NIH grants 1DP2GM119419 and 5U24AI118672. G.D.V. is a Searle Scholar, a Burroughs Wellcome Investigator in the Pathogenesis of Infectious Disease, a Pew-Stewart Scholar, and a MacArthur Fellow.

## Materials and Methods

### Mice

C57BL6/J and B6.SJL, mice were purchased from Jackson Labs (Strain numbers 000664, 002014, and 022486, respectively). PA-GFP (Victora et al., 2010), B1-8^hi^ (Shih et al., 2002), *Ly75*^−/−^ (Inaba et al., 1995), and *Ccnd3*^−/−^ (Kozar et al., 2004), mice were bred and maintained in our laboratory. Some of the B1-8^hi^ *Ly75*^−/−^ mice used in scRNA-seq analysis also carried a FUCCI-Green (Fluorescent Ubiquitination-based Cell Cycle Indicator) allele (Sakaue-Sawano et al., 2008), which was not used in the analysis (See Table S1). *Ccnd3*^T283A^ mice were generated in C57BL6 x SJL F1 zygotes and backcrossed onto a C57BL6 background throughout the duration of the study and all experiments shown use backcross generations N1 to N8. Backcross generations used in this study is as follows: N8 for B1-8^hi^ cell transfer (Fig. 6K-M), N4-6 for early S phase entry (Fig. 6B-G), N6 for scRNA-seq analysis (Fig. 6N-Q), N1 for the longitudinal survival curve (Fig. 7A), N1 or N5 for clonal expansion and tumor formation analysis (Figs. 7B-F and S5). Mice were *H2*s/b for the longitudinal survival curve, and *H2*b/b in all other experiments. Littermate mice were used as controls in all cases. For direct competition assays (Fig. 5A, 6K, S3M), we alternately label wildtype and mutant cells using a PA-GFP transgene or CD45.1/2 congenic markers in order to avoid any technical bias that could be introduced with a given transgene. All mice were housed in groups of 2-5 animals per cage in specific pathogen-free facilities at the Rockefeller University. All protocols were approved by the Rockefeller University Institutional Animal Care and Use Committee. 6-8 week-old male and female mice were used in all experiments, except for adoptive transfers, when all recipient mice were males.

### Cell Transfers, Immunizations, and Treatments

Spleens were dissected from mice of indicated genotype, mashed, and filtered through 40 μm cell strainer, red blood cells were lysed with ACK buffer (Lonza), and resulting cell suspensions were filtered into PBS supplemented with 0.5% BSA and 2 mM EDTA (PBE). Naive B cells were isolated by negative selection with magnetic cell separation (MACS) using anti-CD43 beads (Miltenyi), according to manufacturer’s manual. Prior to cell transfer, percentage of NP-binding B1-8^hi^ cell were determined by staining a fraction of cells with 5 mg/mL NP(19)-PE (Biosearch Technologies) and analyzing by flow cytometry.

To induce GCs, where indicated, male C57BL/6 recipient mice were first primed intraperitoneally (i.p.) with 50 mg ovalbumin (OVA) in alum (Thermo Scientific) at 2:1 v:v ratio in 100 μL total volume, two to four weeks prior to cell transfer. Isolated B1-8^hi^ B cells were adoptively transferred at the indicated proportions as depicted in experimental setup of each figure. One day after adoptive cell transfer, mice were immunized with 10 μg NP(19)-OVA (NP-OVA, Biosearch Technologies) absorbed in alum or 25 μg NP-OVA, for primary or boosting, respectively, by a subcutaneous (s.c.) injection into the hind footpad. Where indicated, 6 days after immunization with NP-OVA, 5 μg of DEC-OVA, produced in our laboratory as previously described (Pasqual et al., 2015), in PBS was injected s.c. into the hind footpad. To block CD40L (Figures 1), mice were injected intravenously (i.v.) with 200 μg of a CD40L blocking antibody (Clone MR-1, BioXCell) or 200 μg of Armenian Hamster IgG isotype control (BioXCell) at the indicated time points. To block MHC Class II (Figure 1), mice were injected i.v. with 200 μg of an MHC Class II (I-A/I-E) blocking antibody (Clone M5/115, BioXCell) or 200 μg of rat IgG2b isotype control (Clone LTF-2, BioXCell) at the indicated time points.

Palbociclib (PD-0332991, SelleckChem) was diluted at 6 mg/ml in PBS and 270 μl were injected i.p., for a total dose of 1.6 mg, or 80 mg/kg.

### Flow Cytometry and Cell Sorting

Cell suspensions were resuspended in PBE and incubated on ice for 30 min with fluorescently-labeled antibodies (Table S2) along with 1 mg/mL of anti-CD16/32 (24G2, eBioscience). For detection of cells in early S of the cell cycle, we performed dual nucleotide pulse and staining as previously described (Gitlin et al., 2014). Briefly, mice were injected i.v. with 1 mg of 5-ethynyl-2-deoxyuridine (EdU) and one hour later with 2 mg of 5-bromo-2-deoxyuridine (BrdU). 30 min after the second injection, lymph nodes were harvested, and single cell suspensions were prepared. After cell surface receptor staining as described above, cells were fixed and permeabilized using BD Cytofix/Cytoperm™ fixation and permeabilization solution and BD Cytoperm Permeabilization Buffer PLUS, respectively. EdU and BrdU incorporation into DNA was assayed using the Click-iT Plus EdU Alexa Fluor 647 Flow Cytometry Assay Kit (Invitrogen) and FITC BrdU Flow Kit (BD), respectively.

For single cell sorting, cells were stained as above and index-sorted directly into 96 well plates containing Buffer TCL (Qiagen) supplemented with 1% β-mercaptoethanol using a BD FACS Aria II. Each plate contained all conditions assayed in each replicate. Cells were washed, filtered, and resuspended in PBE prior to analysis or sorting on BD FACS LSR II, FACS Symphony or FACS ARIA II cytometers. All data were analyzed using Flowjo software v.10.

### Library preparation for single cell RNA-sequencing

Libraries for scRNA-sequencing were prepared as previously described (Trombetta et al., 2014). Briefly, nucleic acids were extracted from sorted single cell using RNAClean XP SPRI beads (Beckman Coulter), and RNA was hybridized first using RT primer (/5BiosG/AAGCAGTGGTATCAACGCAGAGTACTTTTTTTTTTTTTTTTTTTTTTTTTTTTT TVN) then reverse-transcribed into cDNA using TSO primer (AAGCAGTGGTATCAAC-GCAGAGTACATrGrGrG) and RT maxima reverse transcriptase (Thermo Scientific). cDNA was amplified using ISPCR primer (AAGCAGTGGTATCAACGCAGAGT) and KAPA HiFi HotStart ReadyMix (Fisher Scientific), cleaned up using RNAClean XP SPRI beads three times, and tag-mented using Nextera XT DNA Library Preparation Kit (Illumina). For each sequencing batch, up to four plates were barcoded at a time with Nextera XT Index Kit v2 Sets A-D (Illumina). Finally, dual-barcoded libraries were pooled and sequenced using Illumina Hiseq2500 (experiment 1)/Nextseq 550 (experiments 2-4) platform.

### Single-cell RNA-Seq analysis

Raw fastq sequence files generated from Smartseq2 libraries were aligned to mouse genome (v. mm10) with the annotated transcriptome (v. gencode M22) using STAR (v. 2.6)(Dobin et al., 2013). Subsequently, genome-mapped BAM files were processed through RSEM (v. 1.3.1)(Li and Dewey, 2011) for gene quantification. The matrix of gene counts was then, used as input for analysis by the R package Seurat (v. 3.1.4.)(Stuart et al., 2019). Next, the dataset was corrected by using the regularized negative binomial regression (SCTransform) implemented by Seurat (Hafemeister and Satija, 2019). To control the dataset for unwanted sources of experimental variation, we fed the mitochondrial gene abundance and the experimental batches information into SCTransform. Additionally, cells containing more than 20% of sequence reads aligned to mitochondrial genes were excluded prior to normalization.

Next, single cells were clustered and gene expression were evaluated with the Seurat workflow. Gene signature scoring was performed by using Seurat’s AddModuleScore function (See Supplemental Spreadsheet 1 for the complete list).

### Immunoblotting

Cells were lysed in ice-cold lysis buffer containing 50mM HEPES (pH 7.4), 40mM NaCl, 2mM EDTA, 1.5mM sodium orthovanadate, 50mMNaF, 10mM pyrophosphate, 10mM glycerophosphate, and 1% Triton X-100, as well as one tablet of EDTA-free complete protease inhibitors (Roche) per 25 ml. Cell lysates were cleared by centrifugation at 15,000 x G for 10 min. Proteins lysates were denatured in sample buffer, boiled for 5 min, resolved by SDS-PAGE, transferred onto PVDF membranes, and were probed with the indicated antibodies.

### Generation of Ccnd2-null and Ccnd3^T283A^ mice

*Ccnd2*-null and *Ccnd3*^T283A^ mice were generated by CRISPR/Cas9-mediated genome targeting at the Whitehead Institute for Biomedical Research and the Rockefeller University, respectively. For *Ccnd2* targeting, cytoplasmic injection of Cas9 mix into fertilized C57BL6 zygotes at the one-cell stage was performed as previously described (Wang et al., 2013; Yang et al., 2013). The Cas9 mix contained Cas9 mRNA and chimeric sgRNA that was in vitro-transcribed from a synthetic dsDNA template (gBlock, Integrated DNA Technologies) using the MEGAshortscript T7 Transcription Kit (Thermo Fisher Scientific) and purified using MEGAclear™ Transcription Clean-Up Kit (Thermo Fisher Scientific). *Ccnd3*^T283A^ mice were generated in C57BL6 x SJL F1 zygotes using the Easi-CRISPR protocol (Quadros et al., 2017). Cas9 mix contained a crRNA/tracRNA complex and a 200 bp repair oligo centered on the introduced mutations, synthesized as a PAGE-purified ssDNA ultramer (all synthesized by Integrated DNA Technologies).

dsDNA template for chimeric sgRNA transcription used for *Ccnd2* (protospacer sequence in capital letters):

cgctgttaatacgactcactatagggACATCCAACCGTACATGCGCgttttagagctagaaa-

tagcaagttaaaataaggctagtccgttatcaacttgaaaaagtggcaccgagtcggtgctttt

crRNA protospacer sequence used for *Ccnd3*:

GTGAATGGCTGTGACATCTG

Repair oligo for *Ccnd3*^T283A^ (differences from original C57BL6 sequence are in uppercase):

gatcgaagctgccctcagggagagcctcagggaagctgctcagacagcccccagcccag-

tgcccaaagccccccggggctctagTagTcaAgggcccagtcagaccagcGctcc-

Tacagatgtcacagccattcacctgtagcttgagacaggccctctcaggccaccaagcagaggaggggcccctgccaccccctccc

### Mathematical modeling

For mathematical modelling to reproduce the percentage of the *Ccnd3*^+/−^ GC B cells during the GC reaction described in vivo, we used a previously reported agent-based model for simulating the GC reaction with the DisseD framework (Meyer-Hermann, 2020), with modifications. Specifically, the in silico simulation was initiated with 50% WT and 50% *Ccnd3*^+/−^ B cells, for which the maximum number of divisions executed in response to Tfh help was fixed to a decreasing series of percentages (80, 76, 72, 71, 60, 50%) of the maximum number allowed to the WT B cells, with the function regulating the number of divisions modified accordingly (Fig. 6I). For each defined percentage, we performed 60 in silico simulations the best model was selected based on the lowest residual sum of squares (RSS, Eq. 1)

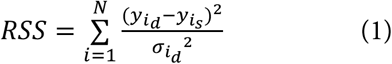

where 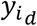 and 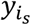 are the experimental and simulated results respectively, 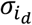 is the standard de-viation of the experimental data and N is the sample size (N = 3).

### Generation of mixed bone marrow chimeras

BM cells were isolated from of punctured tibiae and femurs dissected from mice of indicated *Ccnd3* genotype, as well as J_H_T mice, by centrifugation at up to 10,000 x G for 10 s followed by lysis of red blood cells using ACK buffer. BM cells isolated from *Ccnd3* were mixed with those from J_H_T mice at 20:80 ratio. Prior to BM cell transfer, recipient mice were irradiated twice at 520 rad 4 hours apart.

### Histological analysis of tissue sections

For immunofluorescence, lymph nodes were dissected and immediately fixed in PBS containing 4% paraformaldehyde and 10% sucrose for 1 hour at 4 °C. Fixed tissues were then incubated in 30% sucrose in PBS overnight at 4 °C, embedded in optimum cutting temperature compound the following day, and sliced into 20μm sections using a Leica Cryostat Microtome. For immunostaining, tissue sections were fixed in ice-cold acetone for 10 min at −20 °C, blocked with Streptavidin/Biotin Blocking Kit (Vector Laboratories), incubated with indicated antibodies diluted in PBS containing 5% BSA and 10% normal goat serum. Sections were mounted in Fluoromount-G (SouthernBiotech) prior to imaging on a Zeiss 700 confocal microscope using a 20x objective with numerical aperture of 0.8. Post-acquisition analysis was performed using ImageJ (NIH; http://rsb.info.nih.gov/ij/).

For hematoxylin/eosin and immunohistochemical stainin, organs were fixed in 4% formaldehyde and embedded in paraffin. Five micron-sections of mouse tissues were deparaffinized and heat antigen-retrieved in citrate buffer pH = 6.4, and endogenous peroxidase (HRP) activity was blocked by treating the sections with 3% hydrogen peroxide in methanol. Indirect immunohistochemistry was performed using biotinylated peanut agglutinin (PNA) followed by avidin– horseradish peroxidase or avidin-AP, and developed by Vector Blue or DAB color substrates (Vector Laboratories; Burlingame, CA USA). Sections were counterstained with hematoxylin.

Slides were scanned using a Zeiss Mirax Slide Scanner and photomicrographs were examined using Aperio eSlide Manager (Leica Biosystems;Wetzlar, Germany) or pictures were acquired using a Zeiss Axioskop imaging microscope.

### Statistical analysis

Pairs of samples were compared using the Mann-Whitney U non-parametric test. Multiple comparisons were carried out using the Kruskall-Wallis non-parametric test with Dunn’s multiple comparison test. Chi-square test was used to compare distributions. P values are reported as non-significant (n.s.) when p ≥ 0.05. All statistical analyses were performed using GraphPad Prism v. 8 software.

## Supplemental Figures and Legends

**Figure S1 (related to Fig. 1).**
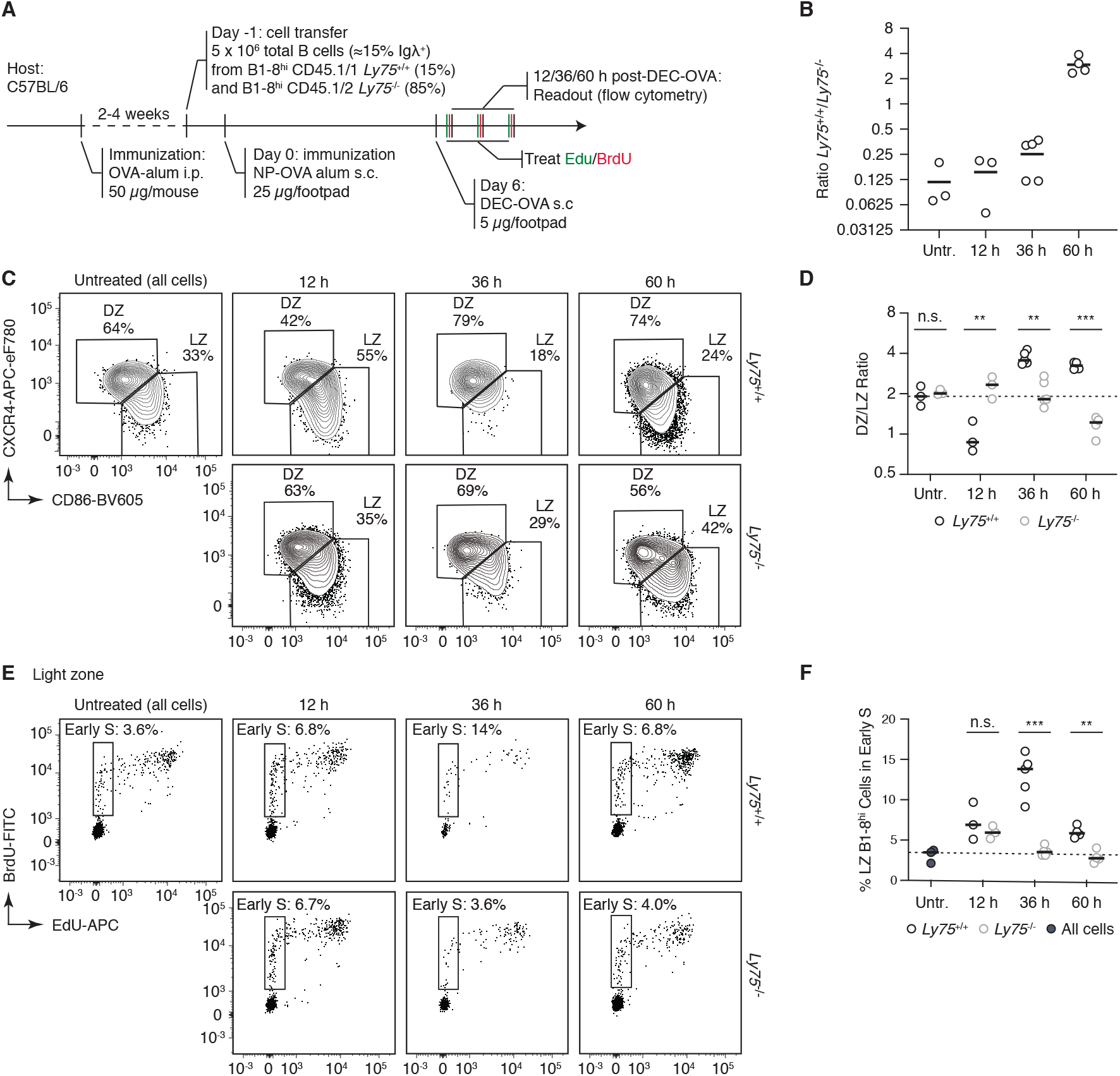
Synchronization and enrichment of *Ly75*^+/+^ cells with DEC-OVA immunization. (A) Experimental setup for DEC-OVA-induced positive selection of *Ly75*^+/+^ GC B cells followed by double labeling with EdU/BrdU to assay for S phase entry. (B) Expansion of *Ly75*^+/+^ cells over *Ly75*^−/−^ over time course. (C) DZ and LZ staining of *Ly75*^+/+^ and *Ly75*^−/−^cells over time course, quantified in (D). (E) S phase entry of *Ly75*^+/+^ and *Ly75*^−/−^ B1-8^hi^ cells in the LZ, quantified in (F). **p < 0.01, ***p < 0.001, paired t test comparison between *Ly75*^+/+^ and *Ly75*^−/−^ cells in the same animal. Bars indicate median. Data pooled from two independent experiments.

**Figure S2 (related to Fig. 2).**
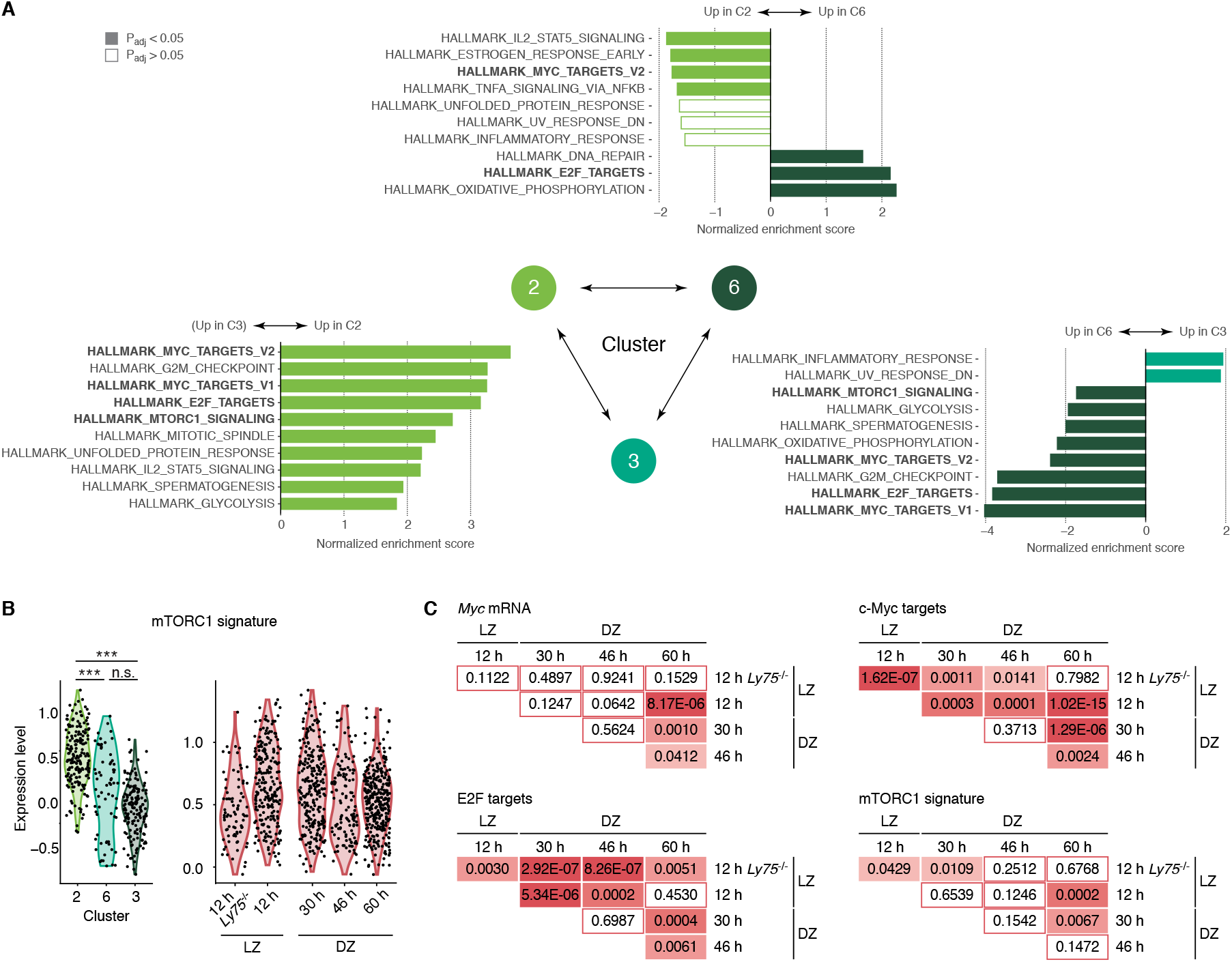
Gene set enrichment analysis (GSEA) between the S phase clusters. (A) Top 10 most significantly varied “hallmark” signatures from the mSigDB database using GSEA to compare clusters 2, 3, and 6. Pathways that are common among three sets of comparisons are indicated in bold. (B) Expression of mTORC1 signature in Clusters 2, 3, and 6 (left) or in DEC-OVA timepoints with indicated zonal phenotypes (right). (C) Summary of p values using Mann Whitney U Test.

**Figure S3 (related to Fig. 3).**
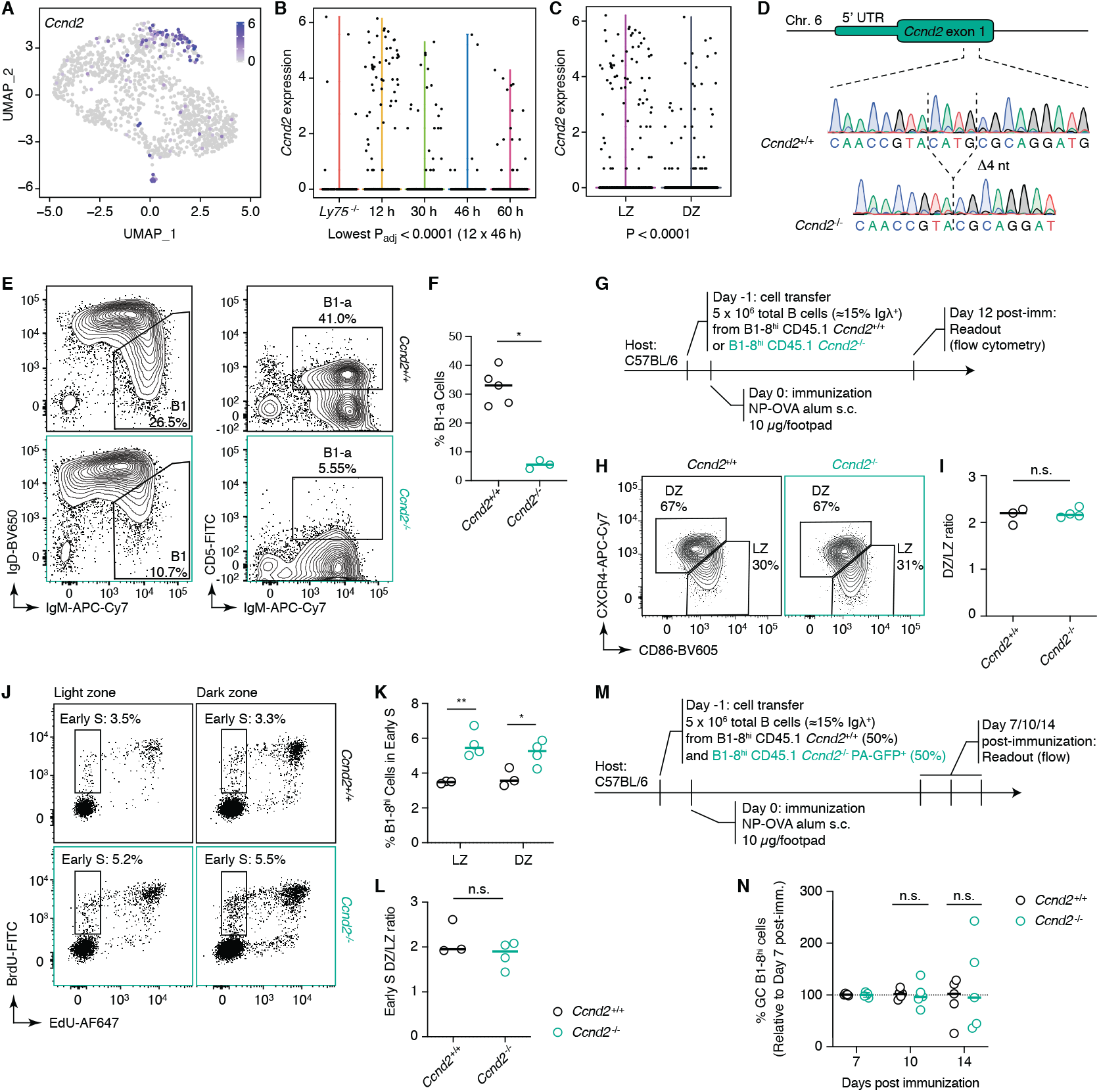
Cyclin D2 is dispensable for inertial B cell cycling. (A-C) Expression of *Ccnd2* in UMAP dimension (A), over time after DEC-OVA immunization (B), and in LZ or DZ (C). P-values in (B) are for Kruskall Wallis test with Dunn’s multiple comparisons test. Other significant P values are: < 0.001 (12 x 60h); 0.0013 (12 vs. 30 h), and 0.037 (*Ly75*^−/−^ vs. 12 h). (D) CRISPR/Cas9-mediated gene targeting strategy to introduce a 4-bp deletion and premature stop codon in *Ccnd2*. (E) Staining for B1 (left) and B1-a (right) cells isolated from peritoneal cavities of WT (*Ccnd2*^+/+^) or cyclin D2 mutant (*Ccnd2*^−/−^), quantified in (F). (G) Experimental setup for induction of GCs containing WT (*Ccnd2*^+/+^) or cyclin D2 mutant (*Ccnd2*^−/−^) B1-8^hi^ cells. (H-L) DZ and LZ staining (H) and S phase entry (J) in WT (*Ccnd2*^+/+^) or cyclin D2 mutant (*Ccnd2*^−/−^) B1-8^hi^ GC cells 12 days after NP-OVA immunization, quantified in (I), (K), and (L). (K) Experimental setup for induction of GCs containing mixtures of WT (*Ccnd2*^+/+^) or cyclin D2 mutant (*Ccnd2*^−/−^) B1-8^hi^ cells. (L) Clonal expansion of WT (*Ccnd2*^+/+^) or cyclin D2 mutant (*Ccnd2*^−/−^) B1-8^hi^ cells over time, relative to Day 7 represented by dotted line. *p < 0.05, **p < 0.01, n.s. non-significant, nonparametric Mann-Whitney test, compared to WT (*Ccnd2*^+/+^) (G, I, and J) or Day 7 (L). Bars indicate median. Data pooled from two independent experiments.

**Figure S4 (related to Fig. 6).**
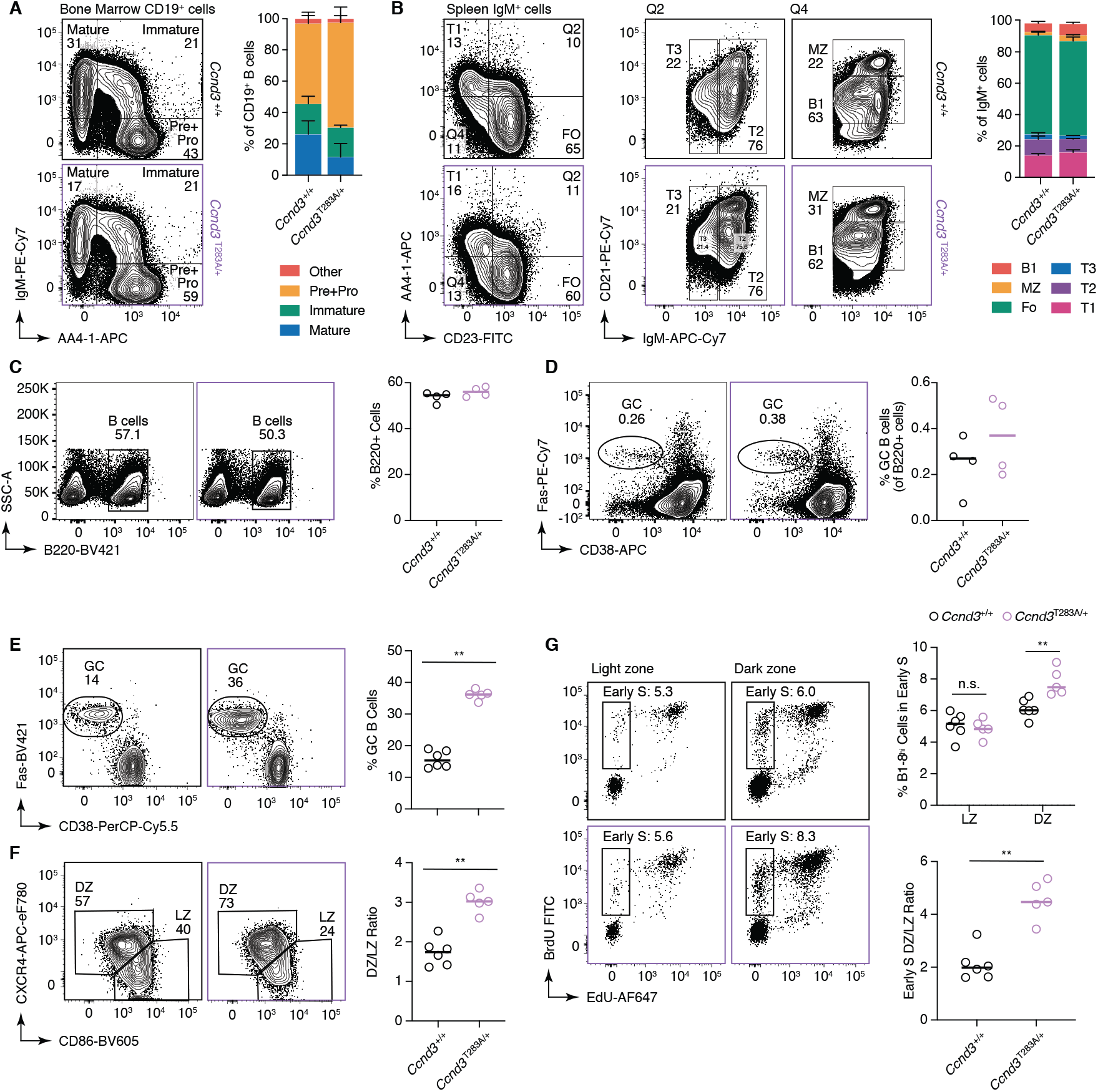
Analysis of *Ccnd3*^*T283A*/+^ mice. (A-B) Bone marrow (A) and splenic (B) B cells development in distribution WT (*Ccnd3*^+/+^) or *Ccnd3*^*T283A*/+^ mice (left) and quantification (right). 6-week-old mice (n = 4 for each genotype) were sacrificed. Bars indicate SD. *Ccnd3*^*T283A*/+^ mice used in this experiment are generation N8. (C-E) Staining (left) and quantification (right)_for GC (C), DZ/LZ (D), and S phase entry of GC B cells induced upon immunization with keyhole limpet hemocyanin (KLH) in pLNs from WT (*Ccnd3*^+/+^) or *Ccnd3*^*T283A*/+^ mice. **p < 0.01, n.s. non-significant, nonparametric Mann-Whitney test. Bars indicate median. Data pooled from two independent experiments. *Ccnd3*^*T283A*/+^ mice used in this experiment are generation N6 and 7.

**Figure S5 (related to Fig. 7).**
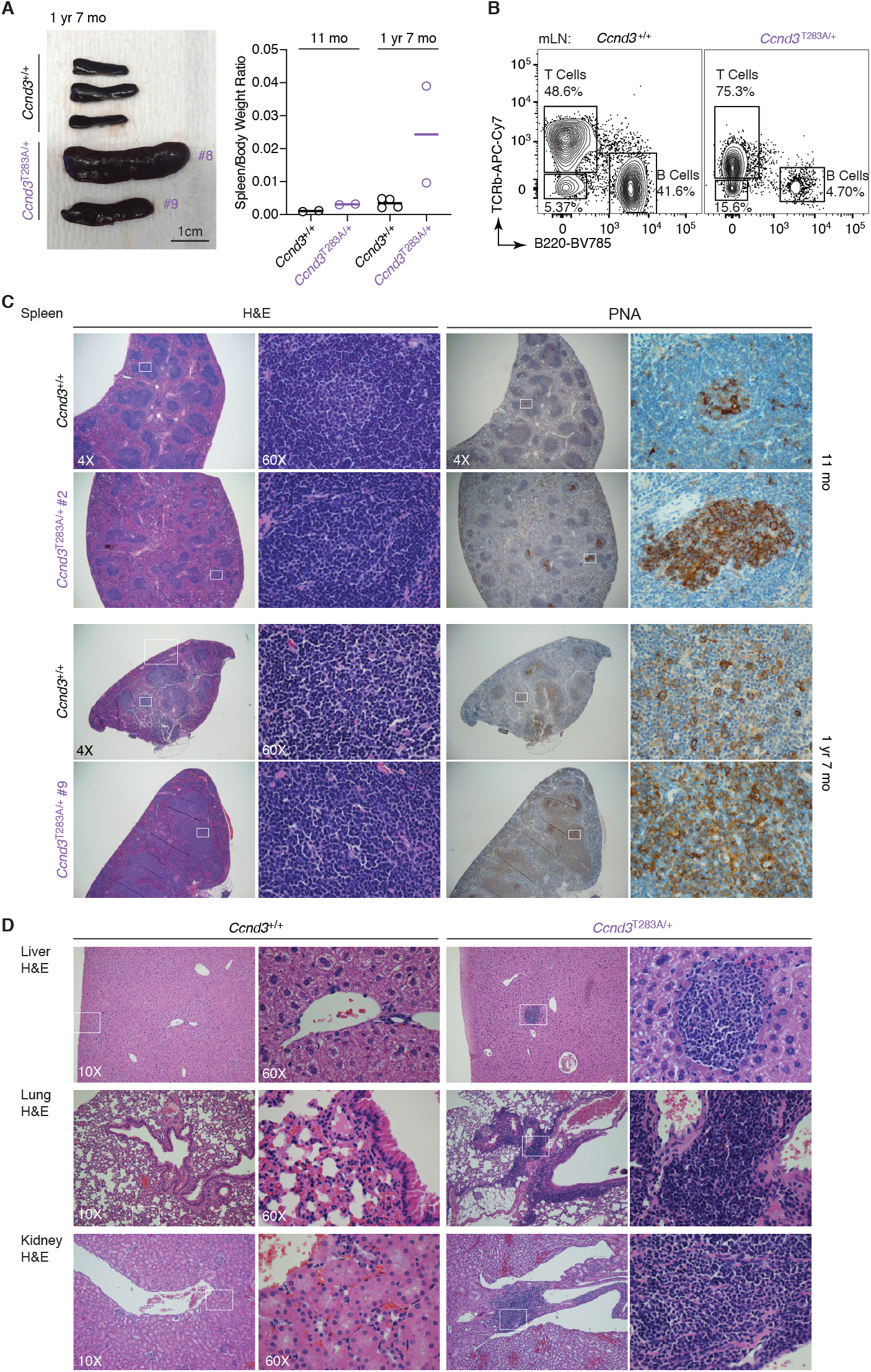
(A) Spleens from WT (*Ccnd3*^+/+^) or *Ccnd3*^*T283A*/+^ animals (right) and quantification of spleen to body weight ratios (left). Bars indicate median. (B) B-T staining in WT (*Ccnd3*^+/+^) or *Ccnd3*^*T283A*/+^ mice revealed expansion of abnormal lymphocyte populations with diminished or no expression of TCRb/B220. (C) H&E and PNA staining of WT (*Ccnd3*^+/+^) or *Ccnd3*^*T283A*/+^ spleens revealed disrupted morphology. (D) H&E of non-lymphoid tissues from WT (*Ccnd3*^+/+^) or *Ccnd3*^*T283A*/+^ revealed lymphocytic infiltration.

**Table S1.**
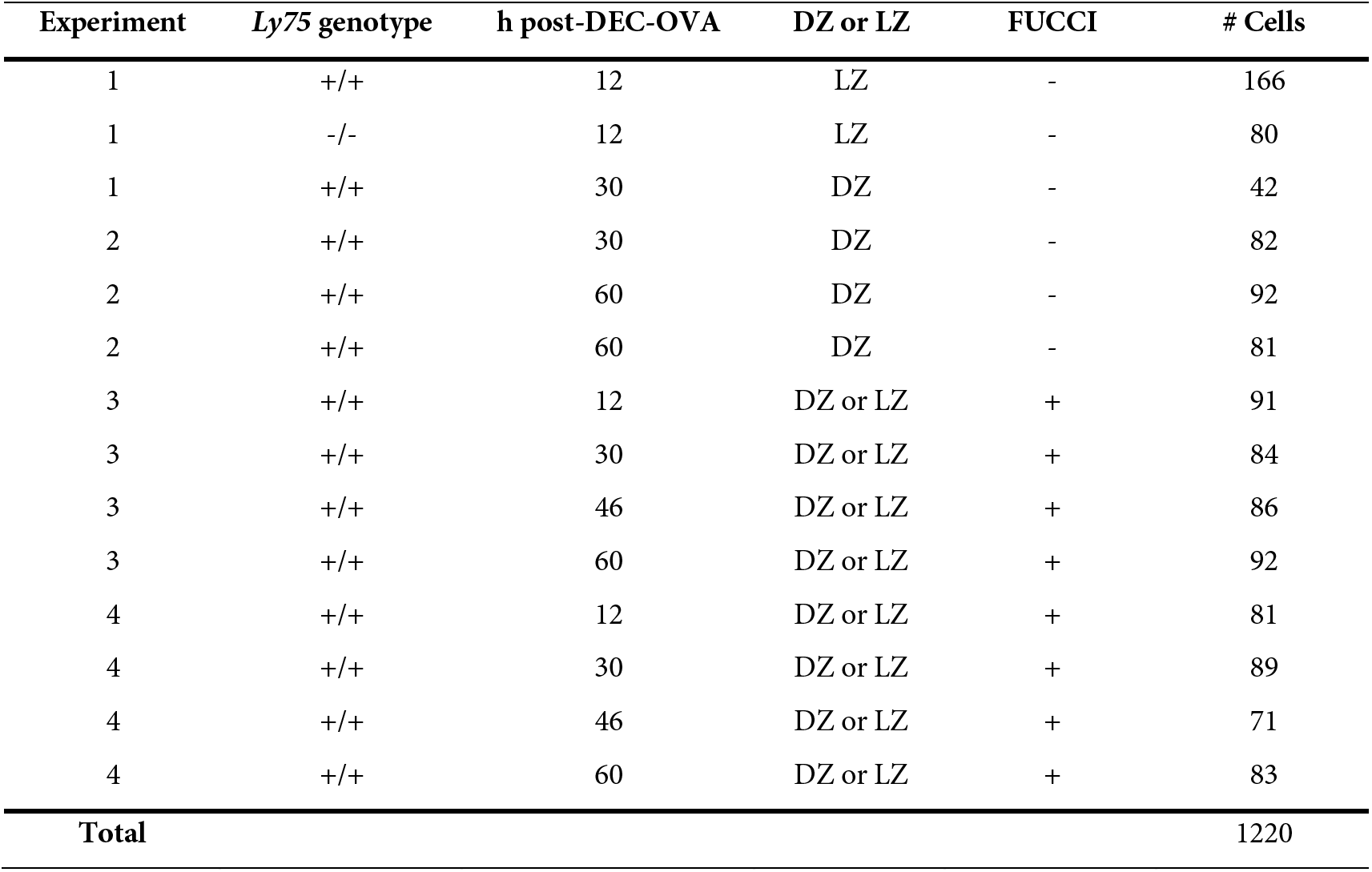
Samples included in single-cell RNA-seq analysis.

| Experiment | *Ly75* genotype | h post-DEC-OVA | DZ or LZ | FUCCI | # Cells |
| --- | --- | --- | --- | --- | --- |
| 1 | +/+ | 12 | LZ | - | 166 |
| 1 | -/- | 12 | LZ | - | 80 |
| 1 | +/+ | 30 | DZ | - | 42 |
| 2 | +/+ | 30 | DZ | - | 82 |
| 2 | +/+ | 60 | DZ | - | 92 |
| 2 | +/+ | 60 | DZ | - | 81 |
| 3 | +/+ | 12 | DZ or LZ | + | 91 |
| 3 | +/+ | 30 | DZ or LZ | + | 84 |
| 3 | +/+ | 46 | DZ or LZ | + | 86 |
| 3 | +/+ | 60 | DZ or LZ | + | 92 |
| 4 | +/+ | 12 | DZ or LZ | + | 81 |
| 4 | +/+ | 30 | DZ or LZ | + | 89 |
| 4 | +/+ | 46 | DZ or LZ | + | 71 |
| 4 | +/+ | 60 | DZ or LZ | + | 83 |
| **Total** |  |  |  |  | 1220 |

**Table S2.** List of antibodies used in the study.

| Conjugate | Antibody | Clone | Source | Application |
| --- | --- | --- | --- | --- |
| BV785 | B220 | RA3-6B2 | BioLegend | FC |
| BV421 | B220 | RA3-6B2 | BioLegend | FC |
| PerCP-Cy5.5 | CD38 | 90/CD38 | BD Pharmingen | FC |
| APC | CD38 | 90 | Invitrogen | FC |
| PE-Cy7 | Fas | Jo2 | BD Pharmingen | FC |
| BV421 | Fas | Jo2 | BD | FC |
| BV711 | CD45.1 | A20 | BioLegend | FC |
| AF700 | CD45.2 | 104 | BioLegend | FC |
| PE | CXCR4 | 2B11 | Invitrogen | FC |
| Biotin | CXCR4 | 2B11 | BD Biosciences | FC |
| Biotin | CD83 | Michel-19 | BioLegend | FC |
| BV650 | CD83 | Michel-19 | BioLegend | FC |
| AF488 | CD86 | GL-1 | BioLegend | FC |
| BV605 | CD86 | GL-1 | BioLegend | FC |
| APC | Biotin | Bio3-18E7 | Miltenyi Biotec | FC |
| APC-eFluor780 | Streptavidin |  | Invitrogen | FC |
| FITC | BrdU |  | BD | FC |
| PE | IgG | RMG1-1 | BioLegend | FC |
| APC | CD138 | 281-2 | BioLegend | FC |
| BV711 | IgD | 11-26c.2a | BioLegend | FC |
| Biotin | CD35 | 8C12 | BD Biosciences | IF |
| AF647 | IgD | 11-26c.2a | BioLegend | IF |
| AF488 | GFP |  | Life Technologies | IF |
| Cy3 | AffiniPure mouse anti-rabbit IgG (H+L) |  | Jackson ImmunoResearch | IF |
|  | Cyclin D3 (mouse) | DCS22 | Cell Signaling | WB |
| HRP | α-Tubulin | 11H10 | Cell Signaling | WB |
| HRP | Mouse IgG1 (goat) |  | Southern Biotech | WB |
|  | MHC class II (I-A/I-E) | M5/114 | BioXcell | In vivo treatment |
|  | Rat IgG2b isotype control, anti-keyhole limpet hemocyanin | LTF-2 | BioXcell | In vivo treatment |
|  | CD40L (CD154) | MR-1 | BioXcell | In vivo treatment |
|  | Armenian hamster IgG | N/A | BioXcell | In vivo treatment |

Flow cytometry (FC), immunofluorescence (IF), western blot (WB)

**Supplemental Spreadsheet 1. List of cell cycle genes, Myc signature genes, E2F target genes, and mTORC1 target genes used in scRNAseq analysis.**

## Notes

### Competing Interest Statement

The authors have declared no competing interest.

